# Long-ranged formation of the Bicoid gradient requires multiple dynamic modes that spatially vary across the embryo

**DOI:** 10.1101/2022.09.28.509874

**Authors:** Thamarailingam Athilingam, Ashwin V.S. Nelanuthala, Catriona Breen, Thorsten Wohland, Timothy E. Saunders

**Affiliations:** Mechanobiology Institute, National University of Singapore, Singapore 117411; Department of Biological Sciences and Centre for Bioimaging Sciences, National University of Singapore, Singapore 117558; Cornell University, Ithaca, NY, USA 14850; Department of Chemistry, National University of Singapore, Singapore 117558; Warwick Medical School, University of Warwick, Coventry, CV4 7AL, United Kingdom

**Keywords:** Fluorescence correlation spectroscopy, Bicoid, *Drosophila* embryo, morphogens, diffusion

## Abstract

Morphogen gradients provide essential positional information to gene networks through their spatially heterogeneous distribution. Yet, how morphogen gradients form is still hotly contested, with multiple models proposed for different systems. Here, we focus on the transcription factor Bicoid (Bcd), a morphogen that forms an exponential gradient across the anterior-posterior (AP) axis of the early *Drosophila* embryo. We utilise fluorescence correlation spectroscopy (FCS) and perturbations to Bcd, to dissect Bcd dynamics at multiple spatial and temporal locations. In both the cytoplasm and nucleus, we find two dynamic modes for Bicoid diffusion dynamics, consisting of fast and slow populations of Bcd. Surprisingly, there are spatial differences in Bcd diffusivity along the AP-axis, with Bcd diffusing more rapidly in the posterior. We establish that such spatially varying differences in the Bcd dynamics are sufficient to explain how Bcd can have a steep exponential gradient in the anterior half of the embryo and yet still have an observable fraction of Bcd near the posterior pole. We subsequently investigated which binding elements of Bcd are playing a role in its dynamics. In the nucleus, we demonstrate that the slower mode of Bcd transport is due to Bcd DNA binding. Addition of the Bcd homeodomain to eGFP::NLS can qualitatively replicate the observed Bcd concentration profile, suggesting this domain is the primary region regulating Bcd dynamics. This study provides a detailed analysis of morphogen dynamics at different spatial and temporal locations, revealing multiple modes of transport. These results explain how a long-ranged gradient can form while retaining a steep profile through much of its range.

## Introduction

Morphogens are signalling molecules that provide crucial spatial and temporal information to cells during development.^1,2^ Knowing how morphogen gradients form and on what time scales is essential for understanding how information can be precisely decoded.^3–6^ Despite intensive study, the underlying dynamics of morphogen gradient formation remain controversial.^7–10^ A longstanding model is that morphogen gradients are formed by localised synthesis and diffusive processes combined with protein degradation or trapping (the SDD model).^3,5,11,12^ Alternative hypotheses for generating a long-ranged gradient include distributed sources of morphogen^13,14^ and long-ranged transport via cytonemes.^15,16^ The shape of the morphogen profile can be adjusted by the mode of degradation,^5,17,18^ which may make the gradient more robust to variation in morphogen protein levels.^17,19,20^ The morphogen profile can also be modulated through receptor binding.^10,21^

A range of quantitative techniques have been used to measure morphogen dynamics *in vivo.* These include fluorescence correlation spectroscopy (FCS),^5,22–24^ fluorescence recovery after photobleaching (FRAP),^3^ single molecule tracking,^25,26^ and protein lifetime measurements.^27–29^ FRAP and protein lifetime measurements provide insight into the longer time dynamics of the system. Essentially, they average out sub-second processes, giving a measure of the effective dynamic parameters across the system. FCS and single molecule imaging have the advantage of measuring the local fast dynamics, but often do not provide information about longer time and spatial processes. See Refs ^7^ and ^18^ for an extended discussion on these points. These differences have led to conflict in the measured dynamic parameters for morphogens. For example, the reported diffusion constant for eGFP::Bcd can vary from 0.4 μm^2^/s (FRAP)^3^ to 7 μm^2^/s (FCS).^30^ While theoretical work has looked to integrate these measurements, taking account of the different time scales measured,^31^ there remains significant contention over the Bcd dynamics. Similar issues are pertinent in other morphogen systems, such as Nodal^24,32,33^ and Wingless.^10,34^ Finally, analysis of morphogen dynamics has typically focused on a fluorescently-tagged version of the wildtype morphogen protein. There is a lack of quantitative data on how morphogen dynamics are altered when protein domains – *e.g.*, DNA or receptor binding motifs - are perturbed. Despite intense study over the past twenty years, it remains a major challenge to dissect the multiple time and spatial scales that underlie morphogen dynamics and the mechanisms that shape the gradient; such knowledge is essential for understanding of how morphogens gradients form *in vivo.*^10^

Here, we take advantage of FCS combined with new reagents for live imaging to dissect the dynamics of Bcd with unprecedented precision. Using this, we reveal, with high accuracy, the dynamic modes that generate the Bcd gradient. We measure the Bcd dynamics at multiple locations along the embryo AP-axis and through nuclear cycles (n.c.) 12-14. We demonstrate that Bcd dynamics do not substantially change over n.c. 12-14, contrary to previous claims.^35^ A two-component fit to the FCS curves is substantially better than a one-component fit in both the cytoplasm and nucleus; this implies two (“slow” and “fast”) dynamic modes in each region. The dynamics of the slower mode correspond closely to measured Bcd dynamics from FRAP.

We show that the effective diffusion coefficient varies across the embryo, with it increasing towards the posterior. These results reveal that Bcd gradient formation is substantially more complicated than previously considered. Implementing such spatial variation within a reaction-diffusion model, we show that the multiple dynamic modes can generate a long-ranged yet steep gradient across a large distance in a relatively short time. Finally, we explored eGFP::Bcd dynamics in a range of mutants with disrupted Bcd binding capacity. Loss of DNA binding increases the fraction of eGFP::Bcd in the fast dynamic mode within the nucleus. In the cytoplasm, we demonstrate that the Bcd homeodomain plays a role in regulating Bcd diffusivity. Combining the Bcd homeodomain with an eGFP::NLS can reproduce the dynamics and gradient shape of the eGFP::Bcd gradient. Overall, we provide a new model – involving spatially dependent dynamics – for formation of the Bcd gradient and we demonstrate that the Bcd dynamics are sensitive to a range of perturbations to binding elements with the protein.

## Results

### Bicoid has multiple dynamic modes and these do not vary across n.c. 12 to 14

Previously, confocal FCS was used to measure the diffusion coefficients of eGFP::Bcd in the anterior cytoplasm^30^ and anterior nuclei.^36^ These measurements were limited in their position and timing within the embryo. We revisited these results, expanding the FCS measurements to both the anterior and posterior regions of the embryo during the interphases of n.c. 12, 13 and 14 (Figure 1A-B, Figure S1A-B). Our FCS autocorrelation curves were highly reproducible between embryos both in the anterior domain and also in the posterior domain where the brightness is very low (Figure S1C). We co-imaged our eGFP::Bcd with H2A::mCherry to ensure that we precisely determined across nuclear cycles (n.c.) and in the cytoplasmic and nuclear regions.

**Figure 1:**
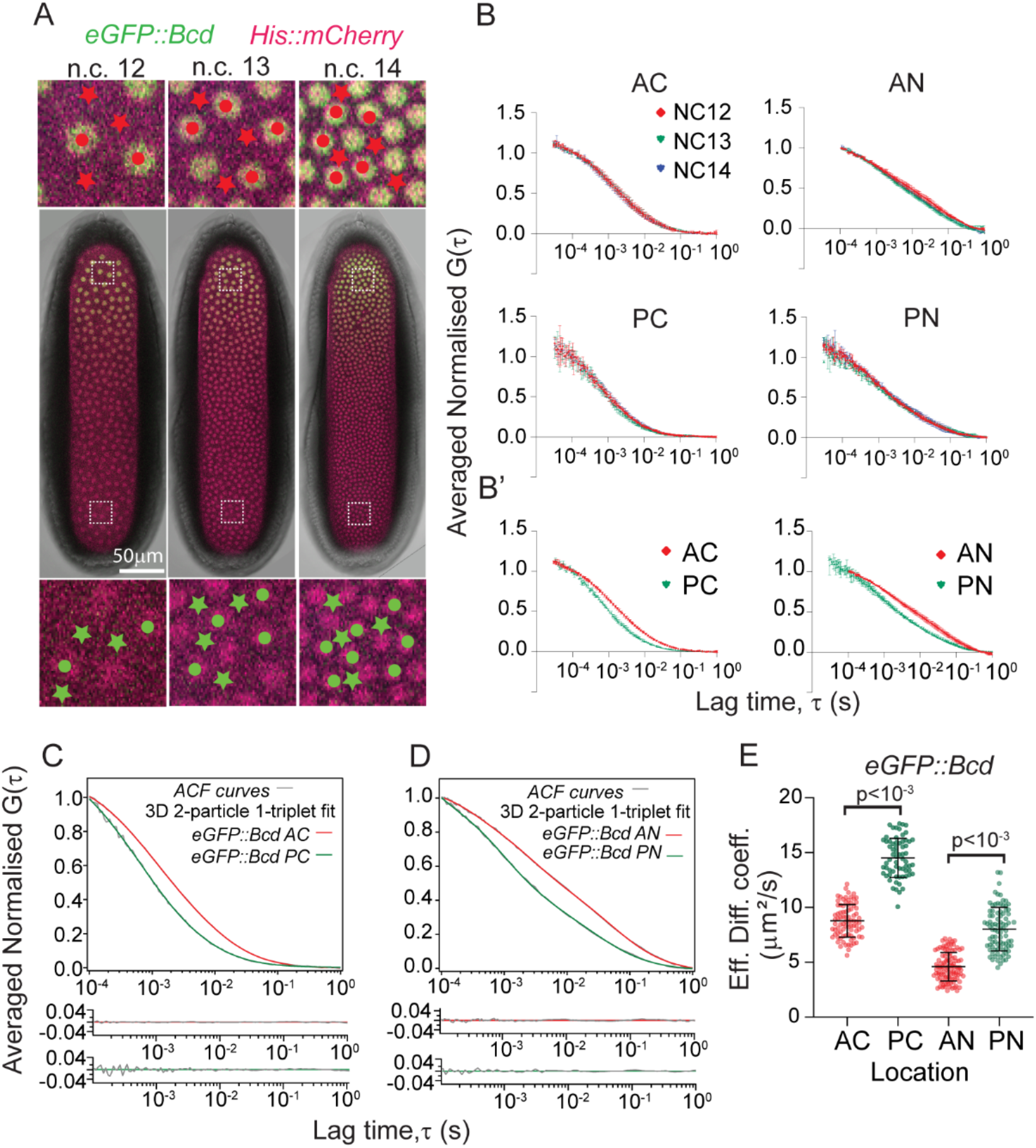
Spatio-temporal dynamics of eGFP::Bcd in the early *Drosophila* embryo. (A) *Drosophila* blastoderm showing the interphase periods of nuclear cycles (n.c.) 12, 13 and 14. Nuclei (mCherry::His2Av, magenta) and eGFP::Bcd (green). Dots and stars indicate cytoplasmic and nuclear regions, respectively, where FCS measurements are carried out in the anterior (red) and posterior (green). (B) Qualitative comparison of normalised and averaged autocorrelation (ACF) curves with mean and S.D. of eGFP::Bcd in the cytoplasmic and nuclear compartments of the n.c. 12,13, and 14 interphases. The anterior and posterior domains are shown in the upper and lower panels respectively. Lag times from 10^-4^ sec to 1 sec are shown. (B’) Superimposed cytoplasmic (left) and nuclear ACF (right) curves from the anterior (red) and posterior (green). (C, D) Comparison of ACF curves (grey) with residues fitted with 3D 2-particle 1-triplet diffusion model. The comparisons are shown from 10^-4^ sec to 1 sec lag times for visual clarity. Red and green fits correspond to anterior and posterior, respectively. (E) Scatter plot comparing the effective diffusion coefficients in different locations within the embryo: anterior cytoplasm (AC), posterior cytoplasm (PC), anterior nuclei (AN), and posterior nuclei (PN). For each condition, multiple measurements are taken from between 3-5 embryos in n.c. 12 to n.c. 14.

Comparing the normalised FCS autocorrelation curves of cytoplasm and nuclei in n.c. 12-14 (Figure 1B, Figure S1C), we saw no clear differences in the curves within or between cycles during interphase. These results reveal that the dynamics of eGFP::Bcd are relatively stable through n.c. 12-14 both at the anterior and posterior locations of the embryo. This contrasts with previous claims based on imaging the Bcd profile within nuclei, which predicted decrease in Bcd diffusion in later stages, D<1μm^2^s^−1^.^35^ We did not explore the dynamics during mitosis with point FCS as the nuclei (chromatin) move rapidly during this phase.

We next fitted dynamic models to the averaged autocorrelation curves (Figure 1C-D). We found that a one-component model of diffusion is insufficient (Figure S1D-E), but a two-component diffusive model (with a fast and slow diffusing population) fitted the data well in all cases (Figure 1C-D). We define the effective diffusion coefficient as *D_eff_* = *f_fast_ D_fast_* + *f_slow_ D_slow_*, where *D_fast_* and *f_fast_* represent the diffusion coefficient and fraction for the fast dynamic mode (and similarly for the slow dynamic mode), with *f_fast_* + *f_slow_* = 1. We found the effective diffusion coefficients of eGFP::Bcd in the anterior cytoplasm and nuclei were 7-9 μm^2^s^-1^ and 4-6 μm^2^s^-1^ respectively (Figure 1E), consistent with previous observations.^30,36^ Therefore, we are confident our results are reproducible and reflect accurate measurements of the eGFP::Bcd interphase dynamics.

### Bicoid dynamics are spatially-dependent in both the nuclei and cytoplasm

We analysed the effective diffusion coefficient in different regions of the embryo. Strikingly, for both nuclear and cytoplasmic eGFP::Bcd, the effective diffusivity was larger in the posterior of the embryo compared with the anterior (Figure 1E). In the posterior, the effective eGFP::Bcd diffusion coefficient in the cytoplasm and nuclei ranged from 12-15 μm^2^s^-1^ and 7-10 μm^2^s^-1^ respectively (Table 1 and 2), ~1.7 fold increase compared to the anterior (Figure S1F). We confirmed this result even if each n.c. was analysed separately (Figure S1G).

Our FCS measurements can be used to estimate the local eGFP::Bcd concentration (Figure 2A) Methods). The measured effective diffusion seems inversely correlated to eGFP::Bcd concentration (compare Figure 1E to Figure 2A). Exploring the different dynamic modes for eGFP::Bcd (Figure 2B-D), we see that in the cytoplasm the slow dynamic mode was similar across the embryo, ~ 1 μm^2^s^−1^ (gure 2?). The fast mode showed an increased diffusivity from *D_fast_* (*anterior*) = 13.0 ± 2.6 μm^2^s ^1^ to *D_fast_* (*posterior*) = 18.8 ± 2.5 μm^2^s ^1^ (Figure 2B, Table 1). In the nuclei, eGFP::Bcd showed a significant, though small, increase in diffusivity in the fast dynamic mode from anterior to posterior regions (9 to 12 μm^2^s^-1^ respectively), with the slow component remaining unchanged (Figure 2B-C, Table 2).

**Figure 2:**
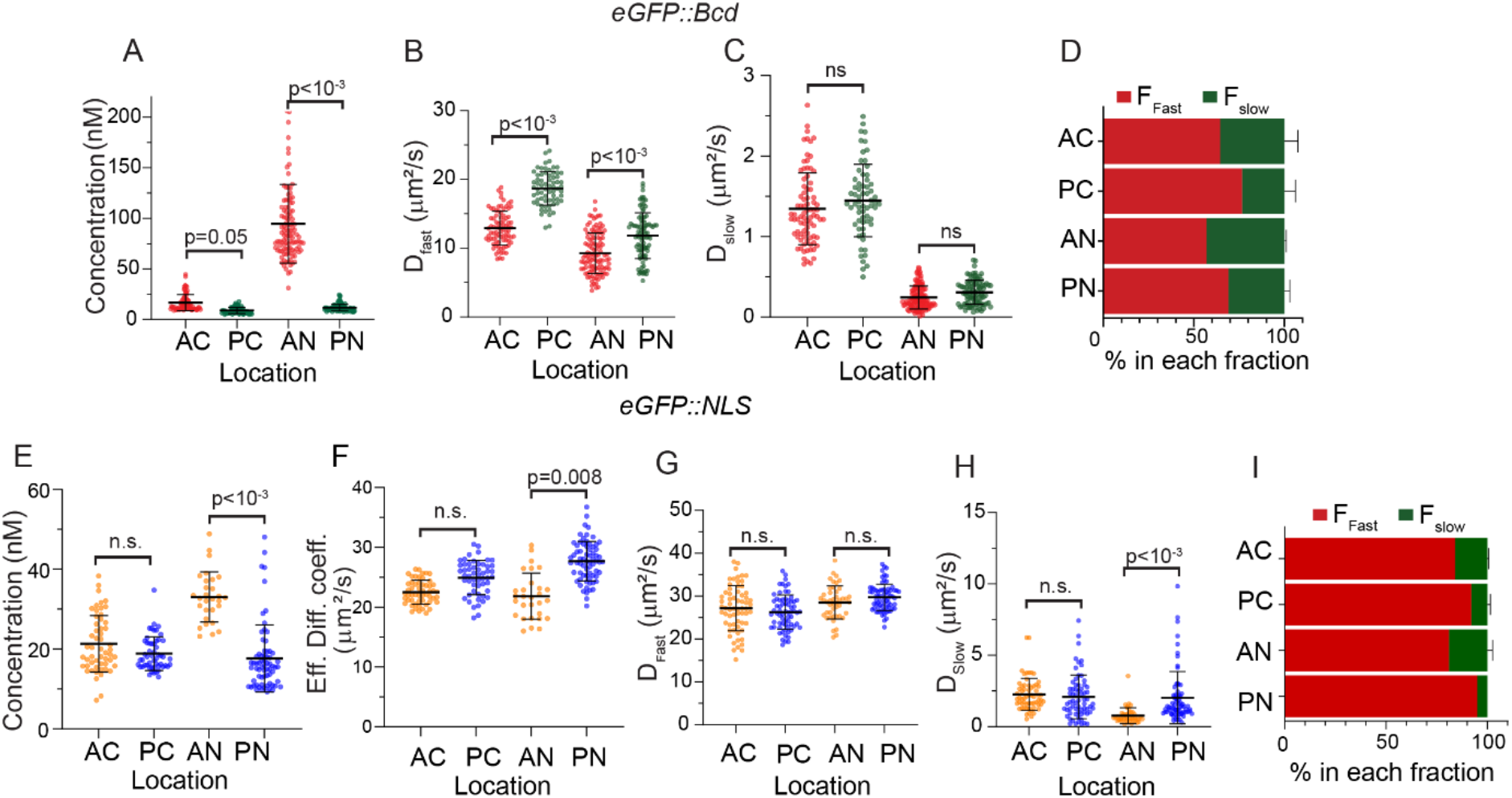
Spatial dependence in Bcd but not NLS dynamics. (A-C) Scatter plot comparing the concentration (A), fast (B) and slow (C) diffusion components of eGFP::Bcd at different locations within the embryo. For each condition, multiple measurements are taken from between 3-5 embryos in n.c. 12 to n.c. 14. (D) Fraction of Bcd in fast and slow dynamic form at different locations within the embryo. (E-I) is same as for (A-D) but for eGFP::NLS, with (F) corresponding to Figure 1E.

We next asked whether the relative fractions of slow and fast components change along the embryo AP-axis. In the anterior cytoplasm, the faster diffusing species comprised *f_fast_* = 65±8 %, whereas in the posterior this increased to *f_fast_* = 78±6 % (Figure 2D, Table 1). Although the slow cytoplasmic dynamic mode was similar in both anterior and posterior of the embryo, the effective diffusion increased towards the posterior because (i) the fast dynamic mode was quicker and (ii) the fraction of the eGFP::Bcd in the fast mode was larger. Likewise, in the nucleus the fraction of eGFP::Bcd in the faster dynamic mode increased towards the posterior, thereby increasing the effective diffusion coefficient (Figure 2D,Table 2).

We performed a similar analysis for eGFP::NLS, driven off the Bcd promoter region. We again fitted with a two-component diffusive model. The majority of the cytoplasmic eGFP::NLS was in the faster diffusive mode (Figure 2E-I, Figure S2, Table 1-2). There was a small difference in the eGFP::NLS dynamics in the nuclei between anterior and posterior ends, possibly due to crowding/non-specific interaction effects in the anterior where protein number is higher (Figure 2E). Supporting this, anterior nuclei show a small but appreciable fraction (~20%) of eGFP::NLS in a slower mode (Figure 2H-I, Table 2). Overall, eGFP::NLS diffusion is more rapid than eGFP::Bcd, with 80-95% existing in the faster diffusive mode. This is consistent with eGFP::NLS only weakly interacting with other components (*e.g.* itself or DNA binding and cytoplasmic elements).

In summary, the dynamics of eGFP::Bcd varies across the embryo, both in terms of the magnitude of the diffusivity and the fraction of slow and fast diffusing eGFP::Bcd. This means that the classic SDD model needs to be revisited as the dynamics are dependent on the spatial location and/or the local morphogen concentration within the embryo.

### Formation of the Bcd gradient with multiple dynamic modes

Given that there exist distinct dynamic modes for Bcd transport across the embryo, we considered the effects of these on long-range gradient formation (Figure 3A). To develop a model of Bcd gradient formation, we need to consider: (1) Bcd has multiple dynamic modes that vary across the embryo; and (2) the observed Bcd gradient is exponential with a decay length λ = 80-100μm across most of the embryo. There are (at least) four effective modes of Bcd transport: (1) fast cytoplasmic fraction; (2) slow cytoplasmic fraction; (3) fast nuclear localised fraction; and (4) slow nuclear localised fraction. We know the fractions in each population in the anterior and posterior of the embryo (Figure 2D). As our focus here is on the long-range establishment of the Bcd morphogen gradient, we considered the nuclear bound fraction to be stationary, and only considered the cytoplasmic component. This assumption is supported by work showing that the Bcd gradient can form with substantially reduced Bcd nuclear import.^37^ To implement the spatially varying diffusion coefficient, we consider 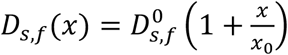 for both slow and fast cytoplasmic components. It is likely that the Bcd diffusivity is really a function of concentration (discussed below), but we use this linear form for simplicity. To account for the changes in the fraction in the slow and fast forms across the embryo, we consider the rate of cytoplasmic Bcd transition from fast to slow forms (denoted by *β*) to behave as 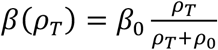 where *ρ_T_* = *ρ_s_* + *ρ_f_* is the total amount of Bcd at each position, *ρ_s_* and *ρ_f_* are the slow and fast forms of Bcd cytoplasmic concentration respectively, and *β*_0_, *ρ_0_* are constants. With this form, the transition rate from fast to slow forms is larger near the anterior pole.

**Figure 3:**
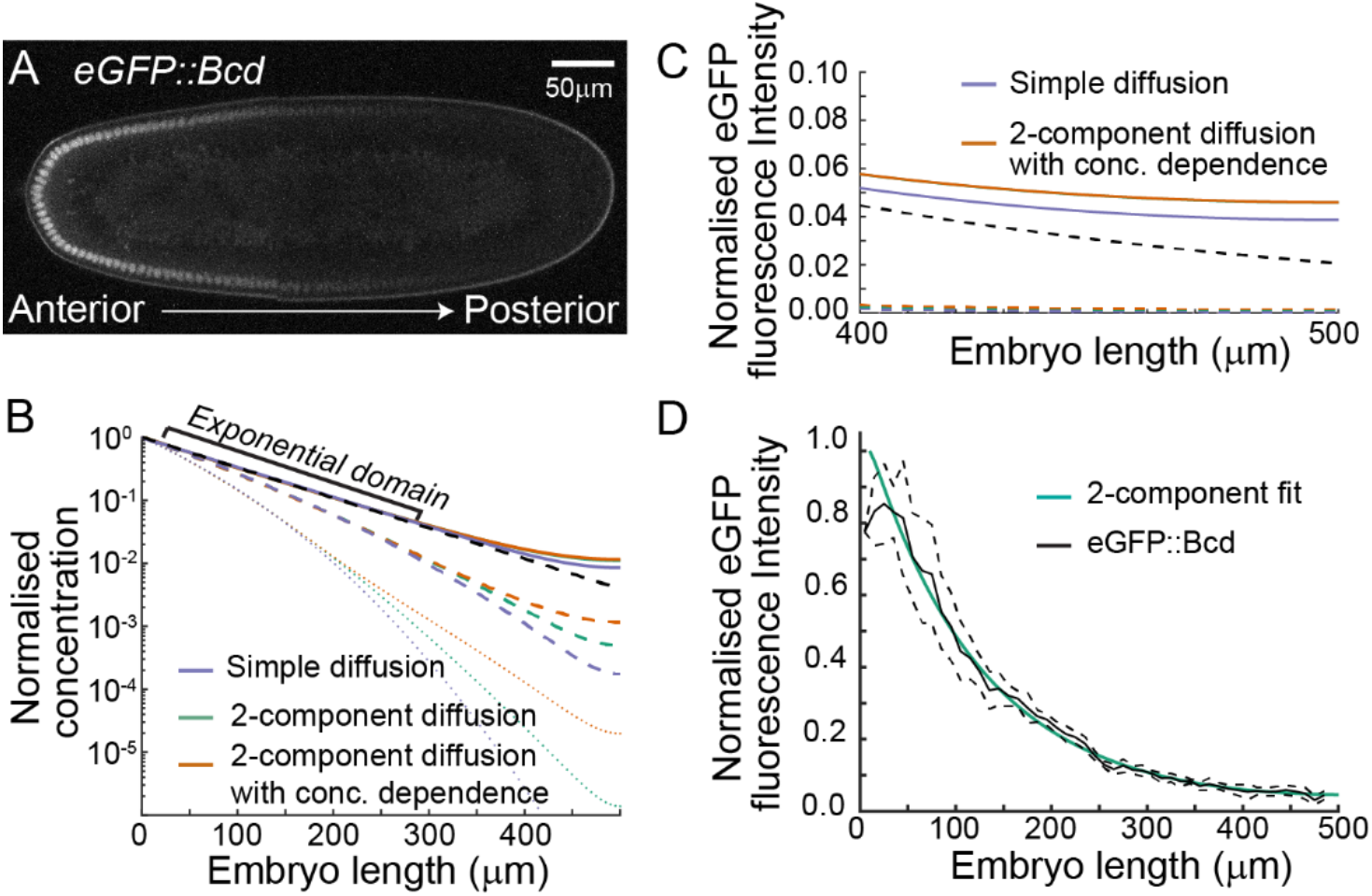
Concentration-dependent parameters within the SDD model can replicate the observed Bcd dynamics and gradient. (A) Embryo expressing eGFP::Bcd, showing the gradient from anterior (left) to posterior (right). (B) Model prediction for formation of the Bcd gradient, where we consider a classic SDD model (blue), 2-component diffusion without parameter sensitivity on concentration (green) and 2-component diffusion with concentration dependence in the switching between fast and slow states (orange). Dotted, dashed and solid lines correspond to 10 min, 25 min and 120 min respectively after initiating the simulation (which starts at zero). (C) Close-up of the predicted concentration profile near the posterior pole in steady-state, showing an increased fraction of Bcd in our 2-component model compared with the SDD model. (D) The 2-component model can accurately fit the observed Bcd concentration profile with parameters based on experimental measurement.

Applying these assumptions leads to the coupled differential equations

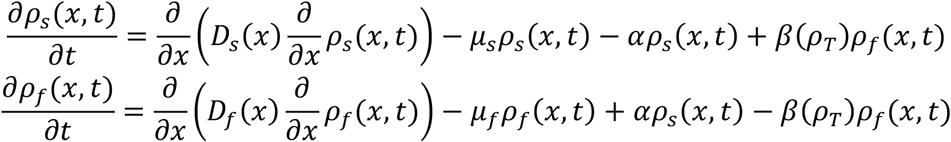

where *μ_s,f_* denotes the degradation rate for the slow and fast forms and *α* is the rate of slow-to-fast transition. For simplicity we keep all parameters constant that are not explicitly highlighted as having a functional dependence. Implementing parameters based on our above and other *in vivo* measurements^28,30,38^ we can plot the Bcd gradient as a function of time and position (Figure 3B).

From this analysis, we see three important points. (1) Having slow and fast forms of Bcd leads to more rapid gradient formation at larger distances (Figure 3B). (2) The increased fraction in the fast form results in increased concentration in the posterior compared to a simple SDD model (Figure 3C), consistent with experimental observation of Bcd in the most posterior.^39^ (3) Even with the multiple species and varying diffusion, the gradient is still exponential across a large extent, consistent with experimental observations (Figure 3D).^3^

In summary, we find that a model of Bcd dynamics that explicitly incorporates fast and slow forms of Bcd (rather than a single “effective” dynamic mode) is consistent with a range of observations that are otherwise incompatible with the standard SDD model. How do these different dynamic modes arise? They are not a simple consequence of spatial differences across the embryo as the eGFP::NLS results show no such spatial dependence. In the next sections, we explore perturbations to binding elements of Bcd to dissect possible mechanisms driving the observed behaviour.

### Bicoid DNA binding determines the slow diffusion dynamics within the nucleus

Given our above observations, we investigated the possible mechanisms regulating Bcd diffusivity. Bcd is a transcription factor known to cooperatively bind to DNA.^40,41^ Further, Bcd binds to *caudal* mRNA, repressing its expression in the cytoplasm. Therefore, we hypothesised that direct DNA/RNA binding impacts Bcd dynamics.

The Bcd homeodomain mutations *bcd^N51A^* leads to loss of downstream target expression (*e.g. hunchback*), posited due to disruption of Bcd DNA binding efficacy.^42^ It also has a reported role in reducing Bcd’s repression of *caudal* expression in the cytoplasm. Another homeodomain mutation *bcd^R54A^* is also reported to de-repress caudal expression but it has normal *hunchback* expression. To test whether these mutations of Bcd alter the protein dynamics, we generated eGFP::Bcd^N51A^ and eGFP::Bcd^R54A^ (Figure 4A). These embryos show a clear anterior-to-posterior gradient of Bcd (Figure 4B).

**Figure 4:**
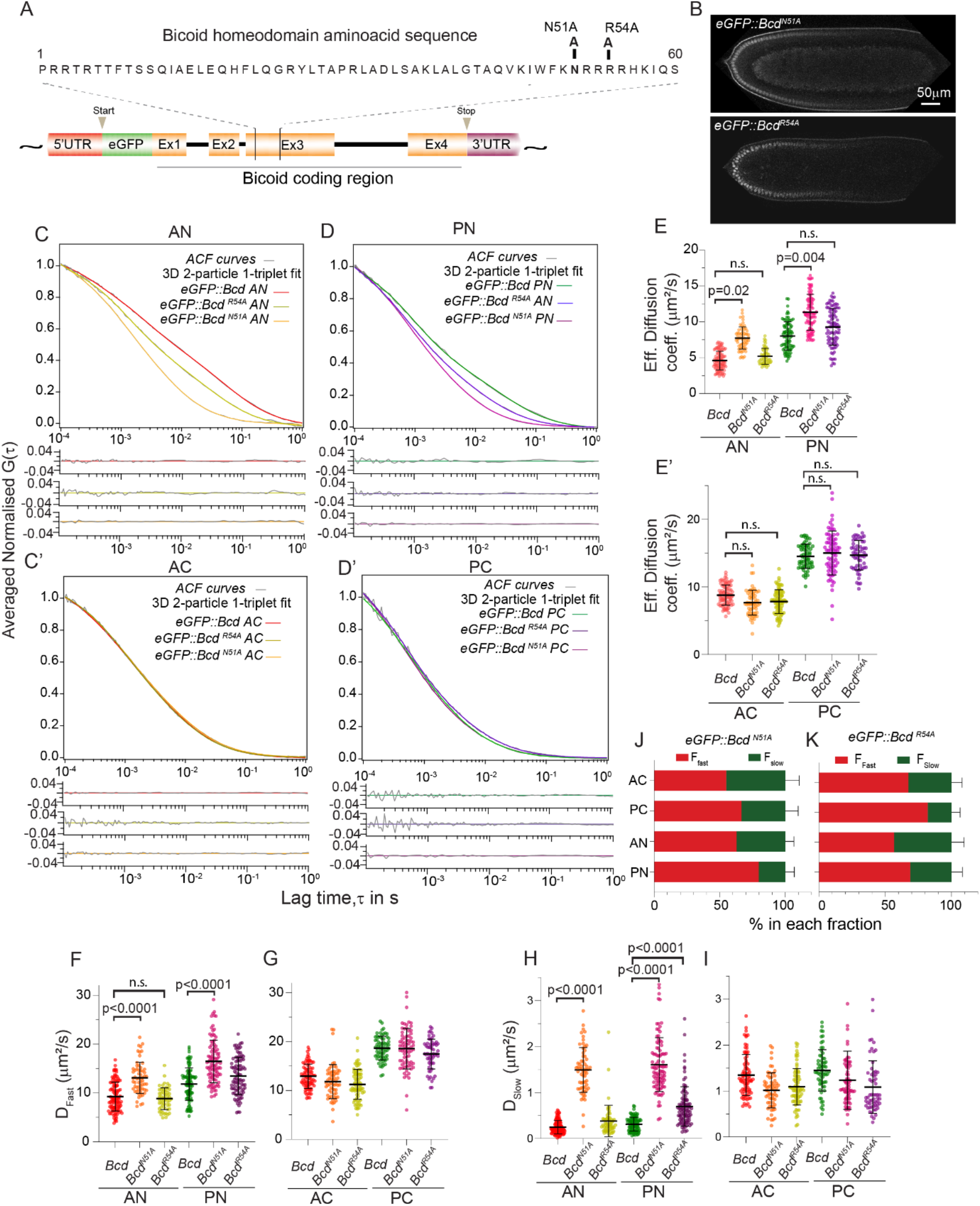
Bcd homeodomain alters nuclear Bcd dynamics. (A) Schematic of *Drosophila bcd* genomic region. 60 amino acids constituting *bcd* homeodomain is expanded. Amino acid conversion at position 51 from asparagine to alanine (N51A) and at position 54 from arginine to alanine (R54A) is displayed. (B) Midsagittal section of n.c. 14 *Drosophila* embryo expressing eGFP::Bcd^N51A^ (top) and eGFP::Bcd^R54A^ (bottom). (C-D’) Comparison of averaged normalised ACF curves (grey) of eGFP::Bcd, eGFP::Bcd^N51A^ and eGFP::Bcd^R54A^ with residues fitted with 3D 2-particle 1-triplet diffusion model. Red and green fits correspond to nuclear and cytoplasmic regions in anterior and posterior regions respectively. Yellow and purple fits correspond to eGFP::Bcd^N51A^. Light green and blue fits correspond to eGFP::Bcd^R54A^. (E-E’) Comparison of the effective diffusion coefficients of eGFP::Bcd and eGFP::Bcd^N51A^ in the nucleus (E) and cytoplasm (E’). (F-I) Scatter plots comparing D_fast_ (H,I) and D_slow_ (J,K) of eGFP::Bcd^N51A^ and eGFP::Bcd^R54A^ at different locations within the embryo (labelling as Figure 1E). (J-K) Bar plots indicating fractions (%) of fast and slow diffusing eGFP::Bcd^N51A^ (J) and eGFP::Bcd^R54A^ (K) molecules in the corresponding embryo compartments. P values calculated using a two-sided permutation test.^43^ All error bars denote ±1 s.d.

We performed FCS on eGFP::Bcd^N51A^ embryos at different locations within n.c. 12-14 embryos (Figure S3). In the nucleus, the normalised autocorrelation curves for eGFP::Bcd^N51A^ embryos were clearly different from eGFP::Bcd embryos, with faster dynamics (Figure 4C-E). However, the dynamics in the cytoplasm were not significantly altered (Figure 4C’-E’). The effective diffusion coefficients further reveal that the diffusion of eGFP::Bcd^N51A^ in the anterior nuclei was 7-9 μm^2^s^−1^ and in the posterior it was 9-13 μm^2^s ^1^ (Figure 4E, Table).

In the nucleus, both the fast and slow modes in eGFP::Bcd^N51A^ embryos show increased diffusivity compared to eGFP::Bcd embryos (Figure 4F-I). While the increase in the slow mode was expected, the reason for the change in the fast mode is less clear. These effects are apparent in both the anterior and posterior of the embryo (Figure S3). As with eGFP::Bcd, the slow mode diffusivity in eGFP::Bcd^N51A^ embryos was similar across the embryo (Figure 4H), with the fast component increasing towards the posterior (Figure 4F). The relative fractions of eGFP::Bcd^N51A^ in the slow and fast modes also changed from anterior to posterior (Figure 4J), with a larger fraction of eGFP::Bcd^N51A^ being in the faster diffusive mode within the posterior nuclei. These results demonstrate that homeodomain function influences the Bcd diffusion dynamics, at least in the nuclei.

The dynamics of eGFP:Bcd^R54A^ embryos were more similar to eGFP::Bcd (Figure 4F-K, Figure S4). There is little difference in the measurements for the diffusion coefficients in nuclei. This is consistent with the R54A mutation not affecting hunchback expression. There is a decrease in *D_slow_* in the posterior, suggesting that the disrupted interaction with *caudal* may be altering posterior Bcd dynamics.

In addition to the homeodomain, the YIRPYL motif and the PEST domain are involved in Bcd-mediated *caudal* mRNA repression. Point mutations in these *bcd* domains are known to abolish the caudal repression in the cytoplasm.^44,45^ Conversions of tyrosine (Y) to alanine (A) and Leucine (L) to Arginine (R) in the YIRPYL motif (YIRPYLàAIRPYR) abolishes caudal repression in the cytoplasm by breaking the interaction of Bcd with the translation initiation factor, eIF4E, at the 5’cap caudal RNA.^45^ Further, replacing 4 threonine (T) and 1 serine (S) residues to 5 alanine (A) residues (Bcd^5aa^) in the PEST domain at positions 188, 193, 195, 197, 200 abolishes caudal repression.^44^ To test the cumulative effect of these binding elements on Bcd dynamics, we generated eGFP:Bcd multi-mutant (MM) line that harbours mutations in the PEST (Bcd^5aa^) domain along with homeodomain N51A and YIRPYL motif (AIRPYR) (Figure S5A). We refer this line as eGFP::Bcd^MM^. FCS measurements of eGFP::Bcd^MM^ embryos in the anterior cytoplasm and nuclei revealed similar Bcd dynamics to eGFP::bcd^N51A^ embryos both in slower and faster Bcd dynamics (Figure S5).

Our results for the N51A, R54A and *bcd^MM^* alleles suggest that: (1) the Bcd homeodomain plays an important role in determining Bcd dynamics; (2) Bcd binding to *caudal* mRNA alone is insufficient to explain the Bcd cytoplasmic dynamics; and (3) there are likely other components (either within Bcd itself or other proteins) in the cytoplasm that affect Bcd dynamics at different Bcd concentrations.

### The Bcd homeodomain regulates protein dynamics in both the nucleus and cytoplasm

Given our above results, combined with the presence of putative cytoplasmic interaction sites within the Bcd homeodomain,^40,41,44,46–48^ we hypothesised that the Bcd homeodomain alone (without the rest of the Bcd protein) may be sufficient to replicate the observed Bcd protein dynamics.

To test this hypothesis, we fused the *bcd* homeodomain to eGFP::NLS, which we refer to as eGFP::NLSbcd^HD^ (Figure 5A). We compared the gradient profiles of eGFP:NLS, eGFP:: NLSbcd^HD^ and eGFP::Bcd (Figure 5B-C). Consistent with our prediction, the gradient of eGFP::NLSbcd^HD^ was steeper than eGFP::NLS (Figure 5C). Remarkably, the eGFP::NLSbcd^HD^ concentration gradient closely matched the eGFP::Bcd profile (Figure 5C). We could also fit this data with a simple SDD model (Methods). This strongly suggests that the homeodomain interactions in the cytoplasm and nuclei are significant contributors to determining Bcd dynamics.

**Figure 5:**
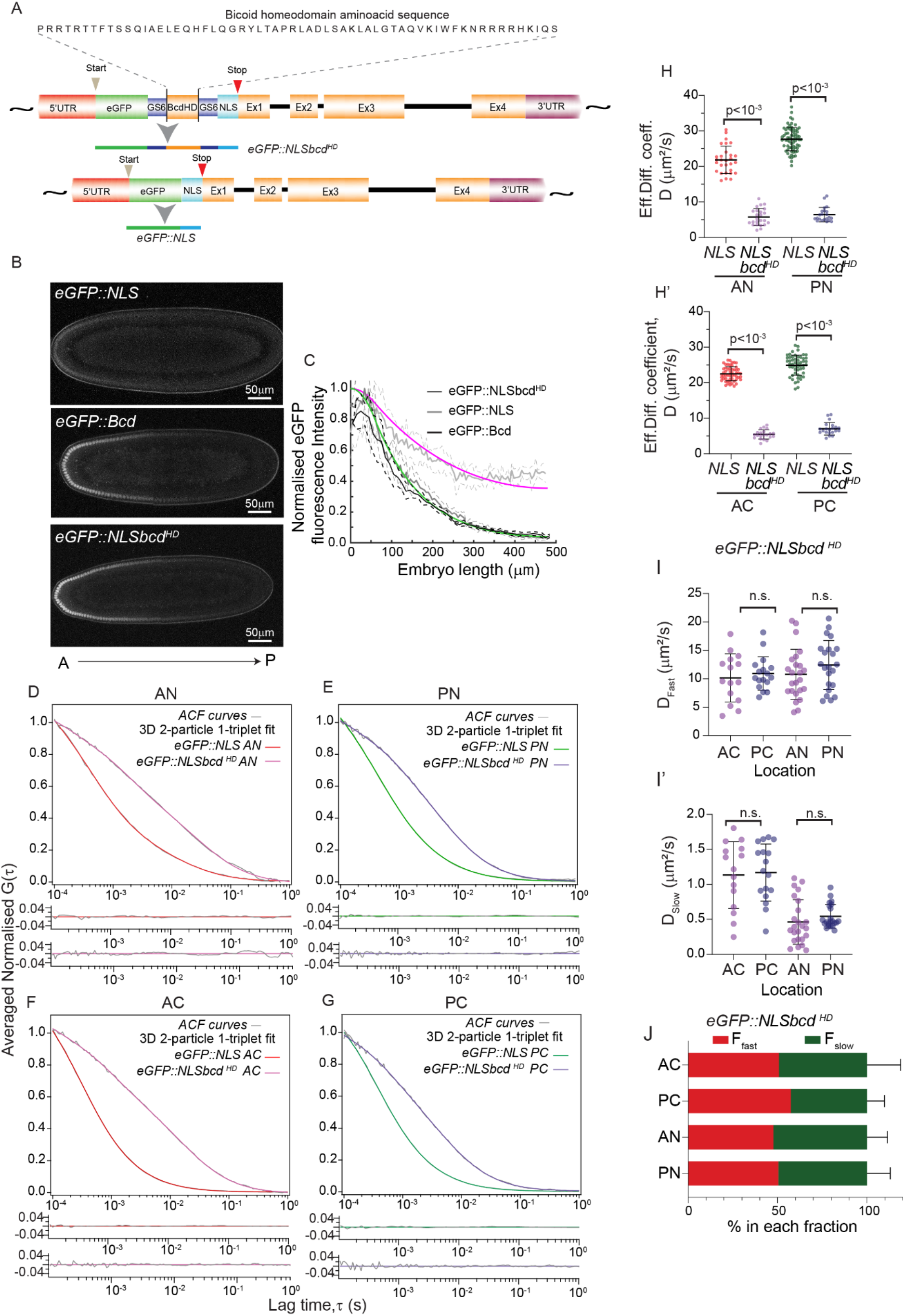
Effects on dynamics of adding the *Bcd* homeodomain to NLS. (A) Schematic of the construction of the eGFP::NLS and eGFP::NLSbcd^HD^ lines. The *bcd* homeodomain is inserted between eGFP and NLS flanked by GS6 linker to generate eGFP::NLSbcd^HD^. (B-C) Midsagittal section of *Drosophila* embryos at n.c.14 expressing eGFP::NLS (top), eGFP::NLSbcd^HD^ (middle), and eGFP::Bcd (bottom). (C) Comparison of normalised gradient profiles between eGFP::NLS (light grey), eGFP::NLSbcd^HD^ (grey) and eGFP::Bcd (black). (D-G) Comparison of averaged normalised ACF curves (grey) of eGFP::NLS and eGFP::NLSbcd^HD^ with residues fitted with 3D 2-particle 1-triplet diffusion model. Red and green fits correspond to nuclear and cytoplasmic regions of anterior and posterior, respectively. Purple and violet fits correspond to eGFP::NLSbcd^HD^. (H-H’) Comparison of the effective diffusion coefficients of eGFP::NLS and eGFP::NLSbcd^HD^ for nuclei (H) and cytoplasm (H’). (I-I’) Scatter plots comparing D_fast_ (I), and D_slow_ (I’) for eGFP::NLSbcd^HD^. (J) Bar plots indicating fractions (%) of fast and slow diffusing eGFP::NLSbcd^HD^ molecules in the corresponding embryo compartments. P values calculated using a two-sided permutation test.^43^ All error bars denote ±1 s.d.

Next, we performed FCS of eGFP::NLSbcd^HD^ embryos to explore the dynamic modes (Figure 5D-G, Table 4). Interestingly, there were no anterior-posterior differences in the diffusion coefficients of eGFP::NLSbcd^HD^ embryos (Figure 5H,H’, Figure S6A’’). This is consistent with the NLS only weakly interacting with other nuclear and cytoplasmic components. This supports the conclusion that the anterior-posterior dynamic changes are due to interactions of the Bcd protein with itself or other elements, and not, for example, due to differences in the physical environment between the anterior and posterior. For eGFP::NLSbcd^HD^, the slow and fast populations represented roughly 50% each at all positions within the embryos (Figure 5J). There is ~3.5-4.5 fold decrease in the effective diffusion coefficients of eGFP::NLS (Figure 5H,H’) upon addition of the homeodomain to NLS encompassing ~2.5 fold decrease in both D_fast_ and D_slow_ values (Figure S6B-E). Further, the fraction of eGFP::NLSbcd^HD^ in the slower form (*F_slow_*) significantly increased compared to eGFP::NLS (compare Figure 2I and Figure 5J). We note that D_slow_ in the nuclei and cytoplasm (0.5μm^2^s^-1^ and 1μm^2^s^-1^) are comparable between eGFP::NLSbcd^HD^ and eGFP:Bcd (0.2-0.3μm^2^s^-1^ and 1μm^2^s^-1^) embryos. This suggests that homeodomain binding affinities to nuclear DNA and cytoplasmic RNA are different and this may regulate the slow Bcd diffusive mode. However, D_fast_ displays distinct differences between eGFP::NLSbcd^HD^ and eGFP:Bcd; the homeodomain is not sufficient to replicate all the Bcd protein dynamics.

## Discussion

We have provided a detailed analysis of Bcd morphogen dynamics in both space and time. Our FCS measurements demonstrate that eGFP::Bcd has both highly motile and slower fractions, in both the nucleus and the cytoplasm. Critically, these dynamics are spatially varying across the embryo. Given the changing fractions of the fast and slow populations in space, the interactions between the populations are likely non-linear, and dependent on the local eGFP::Bcd concentration.

These results suggest that (i) Bcd DNA binding plays an important role in determining Bcd dynamics within the nucleus and (ii) the dynamics of Bcd within the nucleus are more complicated than a simple model of bound versus unbound Bcd.^22^ This might point towards anomalous diffusion as the dominant diffusive mode,^49^ in which instead of two distinct diffusion components, the diffusion coefficient is scale dependent. In fact, the anomaly parameter can provide a good empirical measure for changes in molecular interactions.^22^ However, the anomalous and two-component fit cannot be easily differentiated and the two-component model provides a simpler model and clearer interpretation of the changes due to binding.

Our results help to resolve several observations regarding eGFP::Bcd dynamics. First, FRAP measurements of eGFP::Bcd in the cytoplasm have reported an effective diffusion coefficient in the range 0.3 – 1 μm^2^s^−1^ A^30^ Our measurements of the cytoplasmic slow fraction (D_slow_~1μm^2^s^-1^) are consistent with this. This suggests that the dynamics of the fast fraction were not captured by previous FRAP measurements. Second, single molecules of eGFP::Bcd have been observed in the embryo posterior.^26^ Even with an effective diffusion coefficient of 7 μm^2^s^−1^, few molecules would be expected at the posterior given the estimated Bcd lifetime (30-50 minutes). However, we show that eGFP::Bcd in its fastest form can move quickly (~18 μm^2^s^−1^), and the fraction of eGFP::Bcd in this form increases at lower concentrations. Therefore, it is possible for a subpopulation of eGFP::Bcd to reach the posterior, while the majority of eGFP::Bcd forms a steep concentration profile (Figure 3D). This is consistent with theoretical predictions, that also postulated that Bcd may have spatially varying dynamics.^31^ Bcd also maintains a similar profile across multiple nuclear cycles.^3^ The presence of a rapidly diffusing pool can (at least partially) help to re-establish the gradient quickly after each division. Finally, it has been proposed that Bcd can be produced throughout the embryo, without need for long-ranged diffusive transport.^14^ Yet, our results suggest that >50% of eGFP::Bcd exists in a rapidly diffusing form *(D >* 5 μm^2^s^−1^), which will “wash out” a locally produced gradient. To summarise, our results provide a mechanism for Bcd to have both slow dynamics (as measured by FRAP), and rapid movement in order to set up the gradient across the embryo in a few hours, largely driven by hindered diffusion.

We have also tried to dissect the specific interaction elements of Bcd that drive its dynamics. In the nucleus, the two populations can be largely (though not completely) explained by Bcd binding to DNA. We have shown in the cytoplasm that the ability to transform the eGFP::NLS gradient into one that qualitatively matches the eGFP::Bcd gradient through the addition of the Bcd homeodomain suggests that this region of Bcd is critical in determining Bcd dynamics. Our eGFP::NLSbcd^HD^ construct does not show concentration/spatial dependence, suggesting that the homeodomain is not wholly sufficient to explain the Bcd dynamics. It is possible that Bcd interacts with cytoplasmic elements, including actin and microtubule structures,^50^ which alter its diffusivity.

Such spatially varying dynamics have been hypothesised previously.^51^ The subcellular gradient of Mex-5 within the *C. elegans* embryo has spatially varying dynamics, due to interactions mediated by polarised distribution of PAR proteins^52^ and a recent study in *Xenopus* extracts has shown that cytoplasmic organisation can alter protein diffusivity.^53^ In single molecule tracking of Nodal, multiple dynamic modes have been observed.^25^ In our case, there are no known significant structural differences in the anterior and posterior ends of the embryo at this stage nor are there gradients of polarity. Our results suggest that the Bcd homeodomain has a role in regulating the protein dynamics in the cytoplasm. Bcd binds to the BRE (Bicoid response element) in the caudal 3’UTR and it also binds to the 5’CAP of the caudal mRNA through its PEST domain via adaptor proteins.^42,44,45,54^ Our results with the N51A and bcd^MM^ lines reveal Bcd binding to caudal is unlikely to have a major impact on the diffusion of Bcd in the cytoplasm. One future test of this result is to measure Bcd dynamics in embryos over- /under-expressing *caudal.* On the other hand, our results with NLSbcd^HD^ show that the *bcd* homeodomain does impact protein dynamics in the cytoplasm. However, the specific domains behind this behaviour remain unclear. There are additional factors, such as Zelda, that may also play a role in spatially varying the effective Bcd dynamics.^38^ Finally, our above results suggest that the transition between the slow and fast cytoplasmic forms may depend non-linearly on the local Bcd concentration. More refined spatial dissection of the dynamics may help illuminate this behaviour more clearly. Such behaviour may enable Bcd to adapt to embryos of variable size^55^. One possible other factor may be cytoplasmic flows, which have recently been demonstrated to play a role in refining the Bcd gradient.^56^

Bcd operates as a morphogen within the *Drosophila* blastocyst. Are our observations potentially relevant for other morphogens, which are typically extracellular ligands? Molecules can be hindered either passively due to micro-geometries of the diffusing environment,^25^ or actively stalled by binding and unbinding of the specific receptors on the cell surfaces or transient binding of interacting proteins.^18^ Recent evidence in Nodal suggests that its transport is akin to hindered diffusion,^25^ resulting in an exponential morphogen distribution. During expansion of *Drosophila* wing imaginal discs, the distribution of Dpp activity can be scaled to the size of the tissue via pentagone,^57^ a feedback regulator of Dpp, and Dpp recycling.^58^ In both the above examples, feedback between the cellular environment (including receptor distribution) and the morphogen dynamics setup the gradient. Our results indicate that Bcd dynamics can be also considered as hindered diffusion in the *Drosophila* blastocyst; a balance between diffusion and interactions with the local region (at least in part mediated by the Bcd homeodomain) generate the effective dynamics that create the Bcd gradient. It seems likely that in the cytoplasm, Bcd transport is hindered by the cytoskeletal structures, which could be pertinent for extracellular morphogens. Therefore, we predict that our key observation – that the effective morphogen dynamics are not constant in space – will be relevant for other morphogen systems.

We have used dual colour imaging to ensure that we are recording accurately either nucleus or cytoplasmic pools. An alternative strategy is to use imaging-FCS.^23^ With this approach, the available timescales are reduced (lowest time resolution about 0.1 ms) but spatial cross-correlation can be explored.^23^ This approach also has the advantage of being able to image throughout mitosis, as spatial movements of the nuclei can be accounted for. It will be interesting to explore how Bcd dynamics change during nuclear division in the blastocyst.

Overall, the combination of new reagents with careful FCS measurements has revealed insights into how a morphogen gradient can form across the required spatial and temporal scales. The apparent concentration-dependent dynamics of Bcd provides a mechanism for how the Bcd gradient can form sufficiently fast while also having slower more local dynamics. Outstanding questions include: (1) what interactions are determining Bcd dynamics in the cytoplasm; (2) is Bcd diffusivity concentration-dependent, and if so how; and (3) whether other morphogens display position/concentration dependent dynamics?

## Acknowledgements

This work was funded by a Singapore MOE Tier 3 grant (MOE2016-T3-1-005) to TW and TES and a Singapore MOE Tier 2 grant (MOE2018-T2-2-138) to TES. TES was also supported by an EMBO Global Investigator award and start-up support from University of Warwick, UK. CB was funded by the Global Fellows Program of the College of Agriculture and Life Sciences, Cornell University, USA.

## Author Contributions

TES initiated the project with proof-of-principle data collected by CB. TA performed all the experiments shown here with assistance from AVSN, under the supervision of TW and TES. TA performed the data analysis with support from TW and TES. TA, TW and TES wrote the first draft of the manuscript, with all authors contributing to the final submitted version.

## Methods

### Generation of fly lines

eGFP::Bcd fly line was a gift from Thomas Gregor. The eGFP::Bcd line was generated by introducing eGFP coding region into the N-terminus of the *bcd* coding region after the start codon in the pCaSpeR7 Bcd plasmid.^60–62^ eGFP tagged Bcd was brought in the background of *bcd^1^* null allele to ensure that it is the only source of Bcd in these embryos. eGFP::Bcd was crossed with His2A::mCherry ^63^ to mark the nuclei of early blastoderm embryos, such that the 560-laser line could act as a reference to mark the nucleus and to differentiate the cytoplasmic region in the syncytium. eGFP::Bcd^N51A^ and eGFP::Bcd^R54A^ mutant lines were generated by introducing point mutations at specific locations in the homeodomain of the eGFP::Bcd pCaSpeR7 construct using primers bearing appropriate base pair changes.

For control, we generated an eGFP::NLS line expressed using the Bcd regulatory sequences.^64^ We introduced a nuclear localisation sequence and a stop codon at the end of the eGFP sequence in the eGFP::Bcd pCasper7 plasmid as described in Ref. ^64^ Upon translation, the eGFP::NLS generated a gradient expression across the A-P axis of the embryo (Figure S2A; Figure 5B). For generating eGFP::NLSbcd^HD^ line, we introduced the homeodomain sequence of Bcd that codes for 60 aminoacids. We PCR amplified the 180 bp homeodomain sequence from exon 3 of the Bcd genomic region using primers that had a GS6 linker and Kpn1 restriction site at their extreme ends. The Kpn1-GS_6_-bcd^HD^-GS_6_ –Kpn1 fragment was digested and inserted into the newly introduced Kpn1 site of the acceptor eGFP::NLS plasmid.

### Preparation of embryos for FCS measurements

eGFP::Bcd; His2A::mCherry embryos at n.c. 9 were dechorionated and mounted in PBS on a coverslip in such a way that the dorsal surface of the embryo struck to the surface of the coverslip and faced the objective. The cortical planar surface of the embryo, which contained the maximum number of in-focus nuclei, was selected for FCS measurements (Figure 1A). His2A::mCherry marked nuclei was used as a reference for the nuclear Bcd FCS measurements, and the area devoid of His2A::mCherry was used for the cytoplasmic Bcd measurements.

### FCS calibration and measaurements

Fluorescence Correlation Spectroscopy (FCS) was carried out using a FV1200 confocal microscope (Olympus) equipped with a time-resolved FCS upgrade kit (PicoQuant) at 25^0^C. The 488 nm pulse wave laser line was used to excite eGFP::Bcd through an UplanSApo 60 X NA 1.2 water immersion objective (Olympus). The laser power was optimised using nuclear eGFP::Bcd at n.c. 14. Laser powers of 2-3μW, which had better signal to noise ratio and minimal photobleaching, were used for FCS measurements both in the nucleus and cytoplasm regions of the anterior and posterior domains of the embryo (Methods Figure 1). The fluorescence emission was passed through a 405/488/543/635 dichroic mirror (Chroma Technology), confocal pinhole of one airy unit (120 μm), and then split using a 50/50 mirror plate. The split emission was detected simultaneously by an avalanche photodiode (SPCM-AQR14; PerkinElmer). Dual detector measurements effectively remove after-pulsing information in the FCS curves. The photon counts from the detector were registered by a TimeHarp 260 time-correlated single photon counting board (PicoQuant) and processed by the SymPhoTime ^65^ software (PicoQuant). The same software was also used to calculate the cross-correlation function.

**Methods Figure 1:**
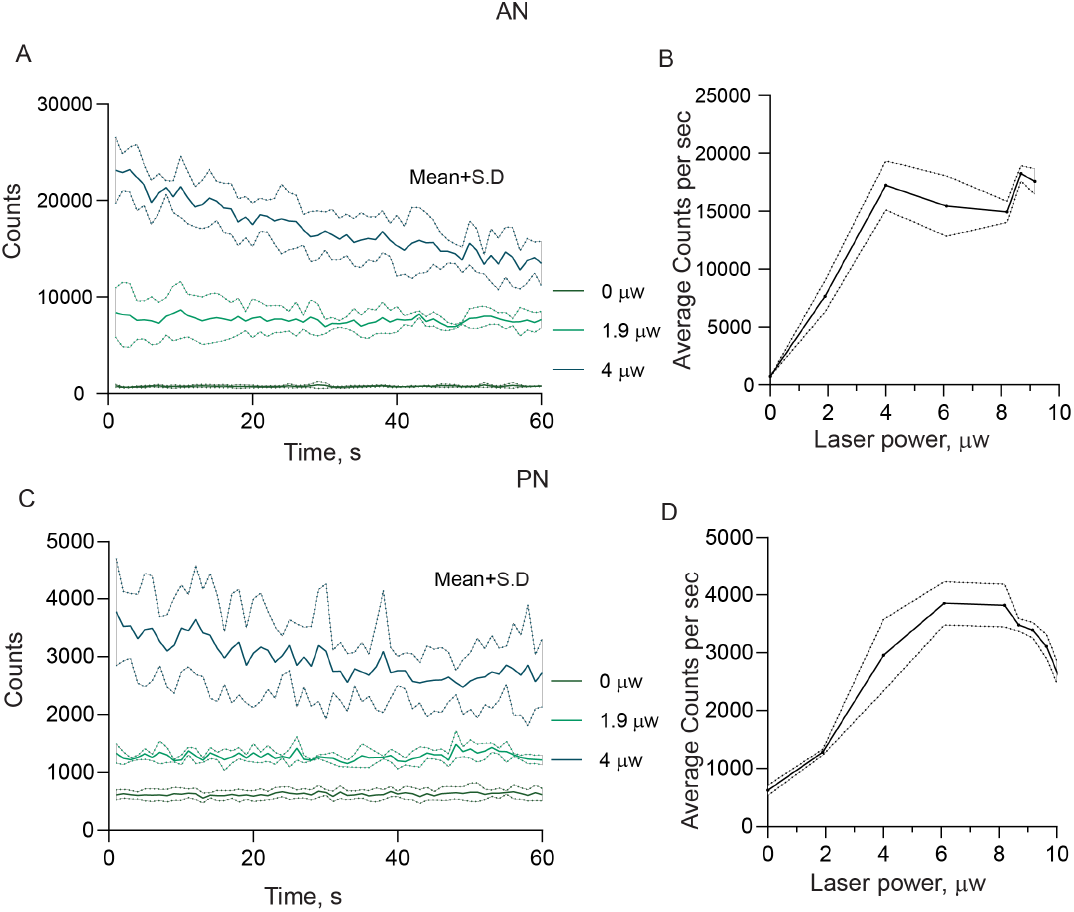
Laser power optimisation. (A, C) line plot showing the time trace of the eGFP:Bcd in the anterior (A) and posterior nuclei (C). Mean (solid line) and SD (dotted line) of the time trace are shown for laser powers 0, 1.9 and 4 μw. (B, D) Mean counts per sec increases with laser power and gets saturated above laser power of 4μw due to photobleaching.

#### Calibration of FCS measurements

We calibrated eccentricity of the confocal volume, k before each round of measurements using the reference dye Atto-488 (Atto-Tec) that has a diffusion coefficient 400 μm^2^/s at room temperature.^65^ The effective volume of calibration (V_eff_, from picoquant) was found to be 0.25±0.04fl and the value of k was found to be 5.6±0.9. The value for k was fixed at 5.6.

#### Estimation of eGFP::Bcd concentration in the measurement volume

We estimated the concentration of eGFP::Bcd and eGFP::NLS through generating a standard curve (Method Figure 2). A high known concentration of Atto-488 dye was serially diluted to lower concentrations (10, 7, 5, and 2 nM) and the correlation amplitudes determined. A plot of the inverse of correlation amplitude *i.e.*, the number of molecules vs concentration in nM fit a linear line (Methods Figure 2B). The unknown concentrations of eGFP::Bcd and eGFP::NLS in the embryos were found from the line equation shown in Methods Figure 2B.

**Method Figure 2:**
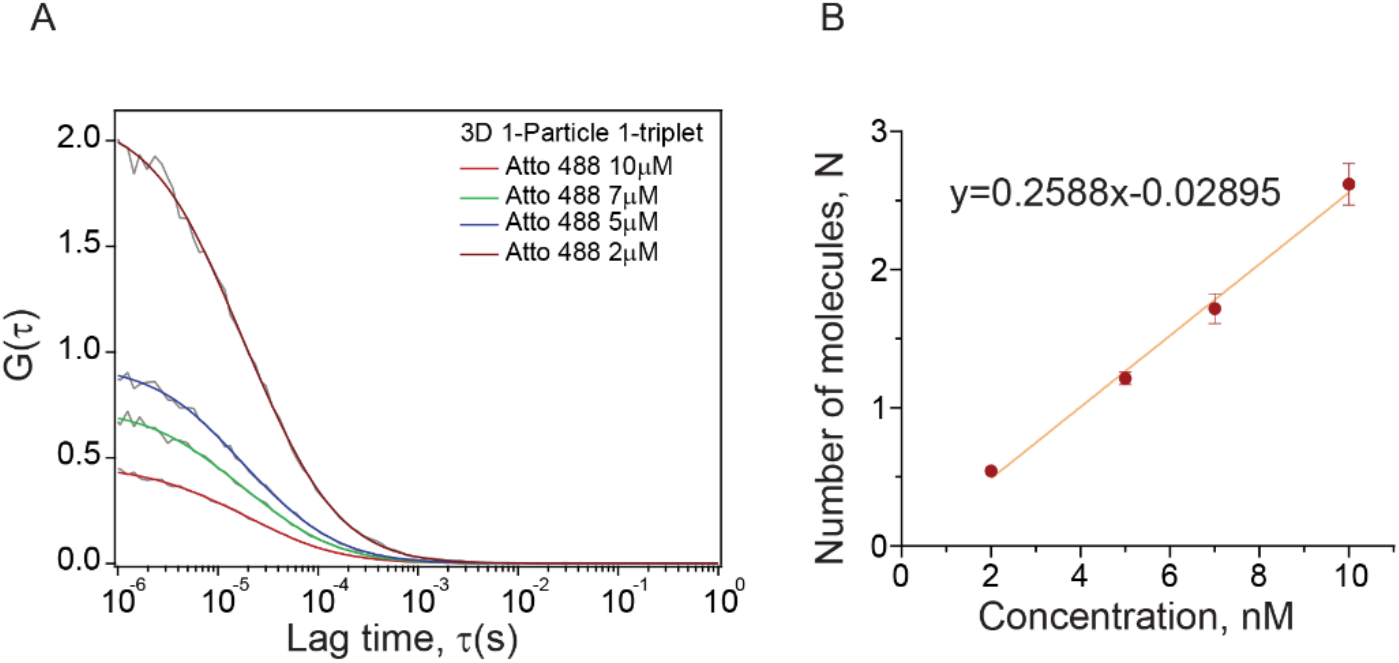
calibration of effective volume to estimate the concentration. (A) Normalised ACF curves (grey) of Atto-488 of different concentration and the 1-particle 1-triplet fits. (B) linear increase in number of molecules in the confocal volume upon increasing the concentration of Atto-488 (2,5,7, and 10nM).

#### Qualitative comparison of ACF curves

ACF curves of individual measurements were normalised and compared to show their qualitative differences. The ACF curves were normalised as *G*(*τ*) – *G*(∞)/*G*(0) – *G*(∞). *G* (0) is the amplitude of the ACF curves (typically at lag time 0.0001s), *G* (∞) is the convergence value at the longer lag times (typically 1s).

### FCS curve Fitting

The FCS curves of each measurement were fitted by 3-dimensional diffusion models involving diffusion of 1 and 2 species with and without reversible switching of the fluorophores to dark states, using Igor-Pro (8.03), FCS data processing plugin, Version 2.1, https://www.dbs.nus.edu.sg/lab/BFL/confocal_FCS.html. The 3D model involving diffusion of 2 species was selected as it provided a good fit determined through quality of the residuals of each plot.^30^ Photophysical processes, *e.g.,* triplet transitions (Atto488) or photoisomerisation and protonation kinetics (eGFP) at short times^66^ are a concern when estimating accurate diffusion times. The error rate in the measurement of the diffusion time becomes worse if the characteristic times of photophysical processes are large enough to overlap with the diffusion time. Further, since the brightness eGFP:Bcd in the anterior and posterior cytoplasm is lower compared to the nuclei, the noise in these curves further limits the distinction between photophysics and diffusion, reducing the accuracy of the determined τ_D_ values.

Therefore, we fitted our FCS curves with different time ranges to include or exclude the photophysical processes at short times. A lag time range of 0.001ms to 1s was considered for fits including photophysics and 0.1 ms to 1 s for fits excluding photophysics. The photophysical parameters, denoted for simplicity as τ_trip_ for the characteristic time and Ftrip for the fraction, were allowed to vary. Comparable τ_D1_ and τ_D2_ values of fits in- or excluding photophysics of each curve were selected. Our estimation reveals comparable photophysics parameters values across the A-P axis (Method Figure 3). The majority of the ***τ_triplet_*** values range from 0.03 to 0.08ms (30 to 80μs) is comparable to the characteristic times of eGFP measured previously.^67,68^ The fraction ranges from 0.15 to 0.22 (Method Figure 3). The data with anomalies due to photobleaching or sudden jumps in the intensities due to movement of the nucleus during measurements, were excluded from evaluation.

**Method Figure 3:**
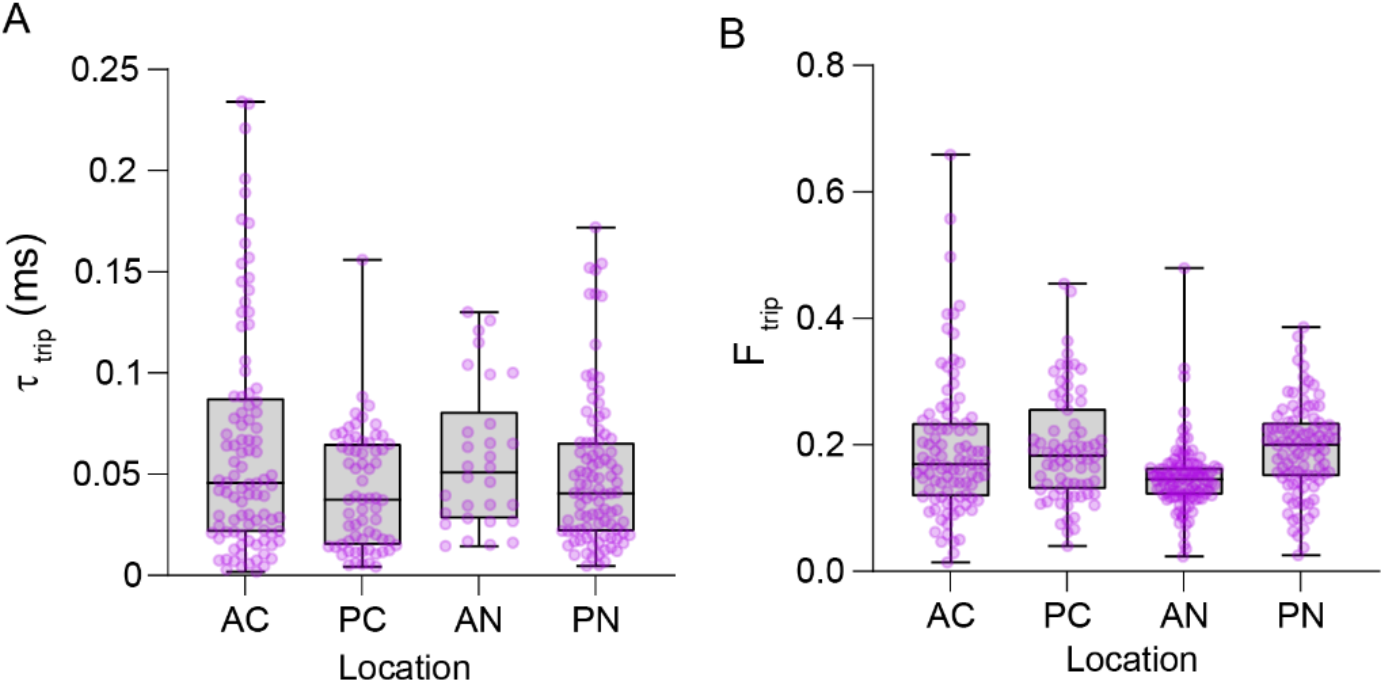
Estimation of triplet lifetime, τ_triplet_ and its fraction (F_triplet_): (A) Box-whisker plot showing the values of the characteristic times of photophysical processes, **τ_triplet_**, from multiple ACF curves measured from 4 embryos, across all compartments. The fraction (Ftrip)is given in B. AC- anterior cytoplasm, PC-posterior cytoplasm, AN-anterior nuclei, and PN-posterior nuclei.

### Quantification of eGFP::Bcd and eGFP::NLS gradients

Time lapse videos of embryos at n.c. 14 were acquired along the longitudinal plane passing through midline of the embryo. For each embryo, two separate images of size 512×512 pixels along the mid sagittal plane covering anterior and posterior domains of the embryo were stitched together. The images were captured at 8bits/pixel, pixel dwell time of 3 μs, line averaging 4, 5 Z-sections of thickness 1 μm each. The conditions captured maximum in-focus peripheral nuclei along the mid-sagittal plane in the Zeiss LSM 710 confocal scanning microscope. The nuclear eGFP intensities along the embryo circumference at n.c. 14 were measured using plot profile plugin of image J. A line mask along the circumference of the embryo was drawn covering the nuclei. The measured raw intensity profile along the circumference of the embryo was background corrected and plotted against the actual length of the embryo.

### Modelling of gradient formation

For the eGFP::NLS model (Figure 5C), we consider the concentration (*ϕ*) as,

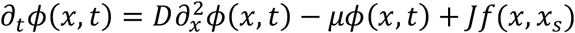

where *D* is the diffusion coefficient, *μ* the degradation rate and *Jf*(*x, x_S_*) defines the production region, and *f*(*x,x_s_*) = 1 *if x* < *x_S_* and *0* otherwise. *D* is taken from the FCS measurements. We normalize out signals, which removes *J* as a free parameter. We take *x_S_* = 20*μm*, consistent with previous observations of *bcd* mRNA. We took *μ* = 1/50*min*. Boundary conditions *∂_x_ϕ*(*x* = 0,*t*) = *0* and *∂_x_ϕ*(*x* = *L,t*) = *0* were used, where *L* = 500*μm* represents the embryo length. For the profiles from eGFP::NLSbcd^HD^ embryos, we used the same parameters as eGFP::NLS, except we reduced the value of *D*, as derived from FCS in Figure 5.

For eGFP::Bcd, the equations are given in the main text. All equations solved in 1D using MATLAB pdepe solver with zero flux boundary conditions at *x*=*L.*

## Data Availability

Raw data of the ACF curves will be provided on request.

## Notes

### Competing Interest Statement

The authors have declared no competing interest.

### Summary of Updates

Reordered material and added new experimental data to strengthen the key message.

## References

1. Briscoe, J. & Small, S. Morphogen rules: design principles of gradient-mediated embryo patterning. Development 142, 3996–4009 (2015).

2. Wartlick, O., Kicheva, A. & González-Gaitán, M. Morphogen Gradient Formation. Cold Spring Harb Perspect Biol 1, a001255 (2009).

3. Gregor, T., Wieschaus, E. F., McGregor, A. P., Bialek, W. & Tank, D. W. Stability and nuclear dynamics of the bicoid morphogen gradient. Cell 130, 141–152 (2007).

4. Kicheva, A. et al. Kinetics of Morphogen Gradient Formation. Science 315, 521–525 (2007).

5. Yu, S. R. et al. Fgf8 morphogen gradient forms by a source-sink mechanism with freely diffusing molecules. Nature 461, 533–536 (2009).

6. Huang, A., Amourda, C., Zhang, S., Tolwinski, N. S. & Saunders, T. E. Decoding temporal interpretation of the morphogen Bicoid in the early Drosophila embryo. eLife 6, e26258 (2017).

7. Huang, A. & Saunders, T. E. A matter of time: Formation and interpretation of the Bicoid morphogen gradient. Curr Top Dev Biol 137, 79–117 (2020).

8. Kerszberg, M. & Wolpert, L. Mechanisms for Positional Signalling by Morphogen Transport: a Theoretical Study. Journal of Theoretical Biology 191, 103–114 (1998).

9. Kornberg, T. B. Cytonemes and the dispersion of morphogens. Wiley Interdiscip Rev Dev Biol 3, 445–463 (2014).

10. Stapornwongkul, K. S. & Vincent, J.-P. Generation of extracellular morphogen gradients: the case for diffusion. Nat Rev Genet 22, 393–411 (2021).

11. Driever, W. & Nüsslein-Volhard, C. A gradient of bicoid protein in Drosophila embryos. Cell 54, 83–93 (1988).

12. Grimm, O., Coppey, M. & Wieschaus, E. Modelling the Bicoid gradient. Development 137, 2253–2264 (2010).

13. Ali-Murthy, Z. & Kornberg, T. B. Bicoid gradient formation and function in the Drosophila pre-syncytial blastoderm. eLife 5, e13222 (2016).

14. Spirov, A. et al. Formation of the bicoid morphogen gradient: an mRNA gradient dictates the protein gradient. Development 136, 605–614 (2009).

15. Roy, S., Huang, H., Liu, S. & Kornberg, T. B. Cytoneme-mediated contact-dependent transport of the Drosophila decapentaplegic signaling protein. Science 343, 1244624 (2014).

16. Stanganello, E. et al. Filopodia-based Wnt transport during vertebrate tissue patterning. Nat Commun 6, 5846 (2015).

17. Eldar, A., Rosin, D., Shilo, B.-Z. & Barkai, N. Self-enhanced ligand degradation underlies robustness of morphogen gradients. Dev Cell 5, 635–646 (2003).

18. Müller, P., Rogers, K. W., Yu, S. R., Brand, M. & Schier, A. F. Morphogen transport. Development 140, 1621–1638 (2013).

19. He, F. et al. Shaping a Morphogen Gradient for Positional Precision. Biophys J 99, 697–707 (2010).

20. Saunders, T. E. & Howard, M. Morphogen profiles can be optimized to buffer against noise. Phys. Rev. E 80, 041902 (2009).

21. Veerapathiran, S. et al. Wnt3 distribution in the zebrafish brain is determined by expression, diffusion and multiple molecular interactions. eLife 9, e59489.

22. Fradin, C. On the importance of protein diffusion in biological systems: The example of the Bicoid morphogen gradient. Biochim Biophys Acta Proteins Proteom 1865, 1676–1686 (2017).

23. Krieger, J. W. et al. Imaging fluorescence (cross-) correlation spectroscopy in live cells and organisms. Nat Protoc 10, 1948–1974 (2015).

24. Wang, Y., Wang, X., Wohland, T. & Sampath, K. Extracellular interactions and ligand degradation shape the nodal morphogen gradient. eLife 5, e13879.

25. Kuhn, T. et al. Single-molecule tracking of Nodal and Lefty in live zebrafish embryos supports hindered diffusion model. Nat Commun 13, 6101 (2022).

26. Mir, M. et al. Dynamic multifactor hubs interact transiently with sites of active transcription in Drosophila embryos. eLife 7, e40497 (2018).

27. Donà, E. et al. Directional tissue migration through a self-generated chemokine gradient. Nature 503, 285–289 (2013).

28. Durrieu, L. et al. Bicoid gradient formation mechanism and dynamics revealed by protein lifetime analysis. Mol Syst Biol 14, e8355 (2018).

29. Khmelinskii, A. et al. Tandem fluorescent protein timers for in vivo analysis of protein dynamics. Nat Biotechnol 30, 708–714 (2012).

30. Abu-Arish, A., Porcher, A., Czerwonka, A., Dostatni, N. & Fradin, C. High Mobility of Bicoid Captured by Fluorescence Correlation Spectroscopy: Implication for the Rapid Establishment of Its Gradient. Biophys J 99, L33–L35 (2010).

31. Sigaut, L., Pearson, J. E., Colman-Lerner, A. & Dawson, S. P. Messages Do Diffuse Faster than Messengers: Reconciling Disparate Estimates of the Morphogen Bicoid Diffusion Coefficient. PLOS Computational Biology 10, e1003629 (2014).

32. Lord, N. D., Carte, A. N., Abitua, P. B. & Schier, A. F. The pattern of nodal morphogen signaling is shaped by co-receptor expression. Elife 10, e54894 (2021).

33. Müller, P. et al. Differential diffusivity of Nodal and Lefty underlies a reaction-diffusion patterning system. Science 336, 721–724 (2012).

34. Stapornwongkul, K. S., de Gennes, M., Cocconi, L., Salbreux, G. & Vincent, J.-P. Patterning and growth control in vivo by an engineered GFP gradient. Science 370, 321–327 (2020).

35. Little, S. C., Tkačik, G., Kneeland, T. B., Wieschaus, E. F. & Gregor, T. The Formation of the Bicoid Morphogen Gradient Requires Protein Movement from Anteriorly Localized mRNA. PLOS Biology 9, e1000596 (2011).

36. Porcher, A. et al. The time to measure positional information: maternal hunchback is required for the synchrony of the Bicoid transcriptional response at the onset of zygotic transcription. Development 137, 2795–2804 (2010).

37. Grimm, O. & Wieschaus, E. The Bicoid gradient is shaped independently of nuclei. Development 137, 2857–2862 (2010).

38. Drocco, J. A., Grimm, O., Tank, D. W. & Wieschaus, E. Measurement and Perturbation of Morphogen Lifetime: Effects on Gradient Shape. Biophys J 101, 1807–1815 (2011).

39. Mir, M. et al. Dense Bicoid hubs accentuate binding along the morphogen gradient. Genes Dev 31, 1784–1794 (2017).

40. Burz, D. S., Rivera-Pomar, R., Jäckle, H. & Hanes, S. D. Cooperative DNA-binding by Bicoid provides a mechanism for threshold-dependent gene activation in the Drosophila embryo. EMBO J 17, 5998–6009 (1998).

41. Lebrecht, D. et al. Bicoid cooperative DNA binding is critical for embryonic patterning in Drosophila. Proc Natl Acad Sci U S A 102, 13176–13181 (2005).

42. Niessing, D. et al. Homeodomain Position 54 Specifies Transcriptional versus Translational Control by Bicoid. Molecular Cell 5, 395–401 (2000).

43. Ho, J., Tumkaya, T., Aryal, S., Choi, H. & Claridge-Chang, A. Moving beyond P values: data analysis with estimation graphics. Nat Methods 16, 565–566 (2019).

44. Niessing, D., Dostatni, N., Jäckle, H. & Rivera-Pomar, R. Sequence interval within the PEST motif of Bicoid is important for translational repression of caudal mRNA in the anterior region of the Drosophila embryo. EMBO J 18, 1966–1973 (1999).

45. Niessing, D., Blanke, S. & Jäckle, H. Bicoid associates with the 5’-cap-bound complex of caudal mRNA and represses translation. Genes Dev 16, 2576–2582 (2002).

46. Ling, J., Umezawa, K. Y., Scott, T. & Small, S. Bicoid-Dependent Activation of the Target Gene hunchback Requires a Two-Motif Sequence Code in a Specific Basal Promoter. Mol Cell 75, 1178–1187.e4 (2019).

47. Ma, X., Yuan, D., Diepold, K., Scarborough, T. & Ma, J. The Drosophila morphogenetic protein Bicoid binds DNA cooperatively. Development 122, 1195–1206 (1996).

48. Yuan, D., Ma, X. & Ma, J. Sequences outside the homeodomain of bicoid are required for protein-protein interaction. J Biol Chem 271, 21660–21665 (1996).

49. Höfling, F. & Franosch, T. Anomalous transport in the crowded world of biological cells. Rep Prog Phys 76, 046602 (2013).

50. Cai, X., Akber, M., Spirov, A. & Baumgartner, S. Cortical movement of Bicoid in early Drosophila embryos is actin- and microtubule-dependent and disagrees with the SDD diffusion model. PLoS One 12, e0185443 (2017).

51. Lipkow, K. & Odde, D. J. Model for Protein Concentration Gradients in the Cytoplasm. Cell Mol Bioeng 1, 84–92 (2008).

52. Griffin, E. E., Odde, D. J. & Seydoux, G. Regulation of the MEX-5 gradient by a spatially segregated kinase/phosphatase cycle. Cell 146, 955–968 (2011).

53. Huang, W. Y. C., Cheng, X. & Ferrell, J. E. Cytoplasmic organization promotes protein diffusion in Xenopus extracts. Nat Commun 13, 5599 (2022).

54. Macdonald, P. M. Translational repression by Bicoid: competition for the cap. Cell 121, 321–322 (2005).

55. Huang, A., Rupprecht, J.-F. & Saunders, T. E. Embryonic geometry underlies phenotypic variation in decanalized conditions. eLife 9, e47380 (2020).

56. López, C. H. et al. Two-fluid dynamics and micron-thin boundary layers shape cytoplasmic flows in early <em>Drosophila</em> embryos. bioRxiv 2023.03.16.532979 (2023) doi:10.1101/2023.03.16.532979.

57. Hamaratoglu, F., de Lachapelle, A. M., Pyrowolakis, G., Bergmann, S. & Affolter, M. Dpp signaling activity requires Pentagone to scale with tissue size in the growing Drosophila wing imaginal disc. PLoS Biol 9, e1001182 (2011).

58. Romanova-Michaelides, M. et al. Morphogen gradient scaling by recycling of intracellular Dpp. Nature 602, 287–293 (2022).

59. Gregor, T., Tank, D. W., Wieschaus, E. F. & Bialek, W. Probing the limits to positional information. Cell 130, 153–164 (2007).

60. Thummel, C. S. & Pirrotta, V. New pCasper P-element vectors. D. I. S. 71,.

61. Hazelrigg, T., Liu, N., Hong, Y. & Wang, S. GFP expression in Drosophila tissues: time requirements for formation of a fluorescent product. Dev Biol 199, 245–249 (1998).

62. Barolo, S., Carver, L. A. & Posakony, J. W. GFP and beta-galactosidase transformation vectors for promoter/enhancer analysis in Drosophila. Biotechniques 29, 726, 728, 730, 732 (2000).

63. Krzic, U., Gunther, S., Saunders, T. E., Streichan, S. J. & Hufnagel, L. Multiview light-sheet microscope for rapid in toto imaging. Nature Methods 9, 730–733 (2012).

64. Gregor, T., McGregor, A. P. & Wieschaus, E. F. Shape and function of the Bicoid morphogen gradient in dipteran species with different sized embryos. Dev Biol 316, 350–358 (2008).

65. Kapusta, P. Absolute Diffusion Coefficients: Compilation of Reference Data for FCS Calibration. Picoquant application note (2010).

66. Widengren, J., Mets, Ü. & Rigler, R. Photodynamic properties of green fluorescent proteins investigated by fluorescence correlation spectroscopy. Chemical Physics 250, 171–186 (1999).

67. Haupts, U., Maiti, S., Schwille, P. & Webb, W. W. Dynamics of fluorescence fluctuations in green fluorescent protein observed by fluorescence correlation spectroscopy. Proc Natl Acad Sci U S A 95, 13573 (1998).

68. Jiménez-Banzo, A., Nonell, S., Hofkens, J. & Flors, C. Singlet Oxygen Photosensitization by EGFP and its Chromophore HBDI. Biophysical Journal 94, 168 (2008).

