## Supplementary Information for "Long-ranged formation of the Bicoid gradient requires multiple dynamic modes that spatially vary across the embryo"

**Contents:**

**Supplementary Figures**

**Supplementary Tables**

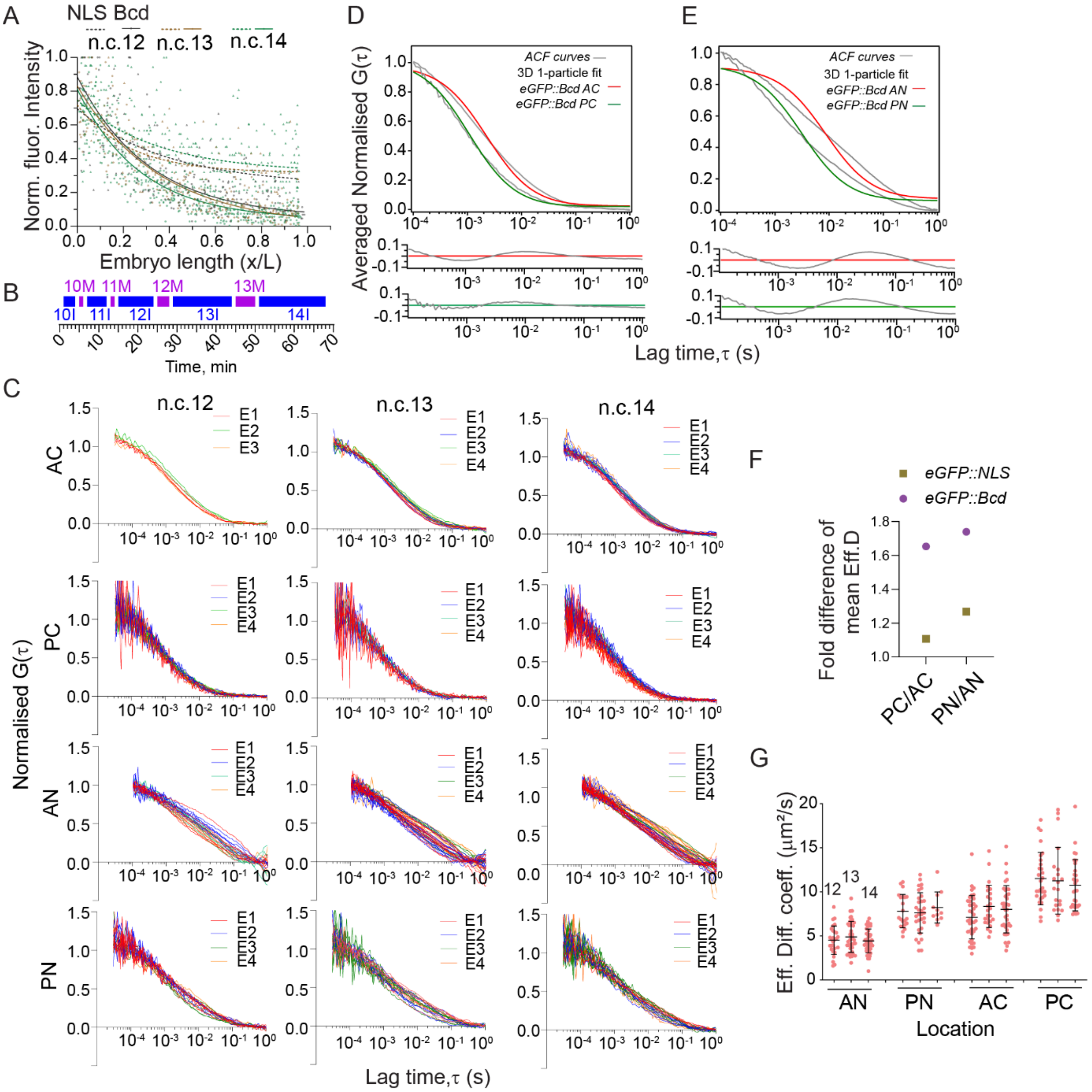


**Figure S1: FCS and fitting of eGFP::Bcd (related to Figure 1)**

(A) Comparison of the gradient profiles of eGFP::Bcd and eGFP::NLS plotted with normalised fluorescence intensities in Y-axis and the normalised embryo length (x/L). Dots show individual nuclear intensities, and the solid lines are fits to exponential profiles. (B) Time profile of the early embryo from n.c. 10 to 14 at 25ºC. I indicates interphase, and M, mitosis. (C) Normalised ACF curves of eGFP::Bcd in the cytoplasmic and nuclear compartments of anterior (Anterior Cytoplasm, AC, Anterior Nuclei, AN) and posterior (Posterior Cytoplasm, PC, Posterior Nuclei, PN) in nuclear cycle (n.c.) 12, 13, and 14 interphases. (D, E) Comparison of ACF curves (grey) with residues fitted using 3D 1-particle diffusion model in the cytoplasm (D) and nucleus (E). (F) Effective diffusion coefficients of nuclear and cytoplasmic locations of the anterior and posterior domains are compared for individual n.c. 12, 13, and 14. (G) Fold change in the mean diffusion coefficient across the embryo for the cytoplasmic and nuclear compartments. Comparison for eGFP::Bcd (circles) and eGFP::NLS (squares) are shown.

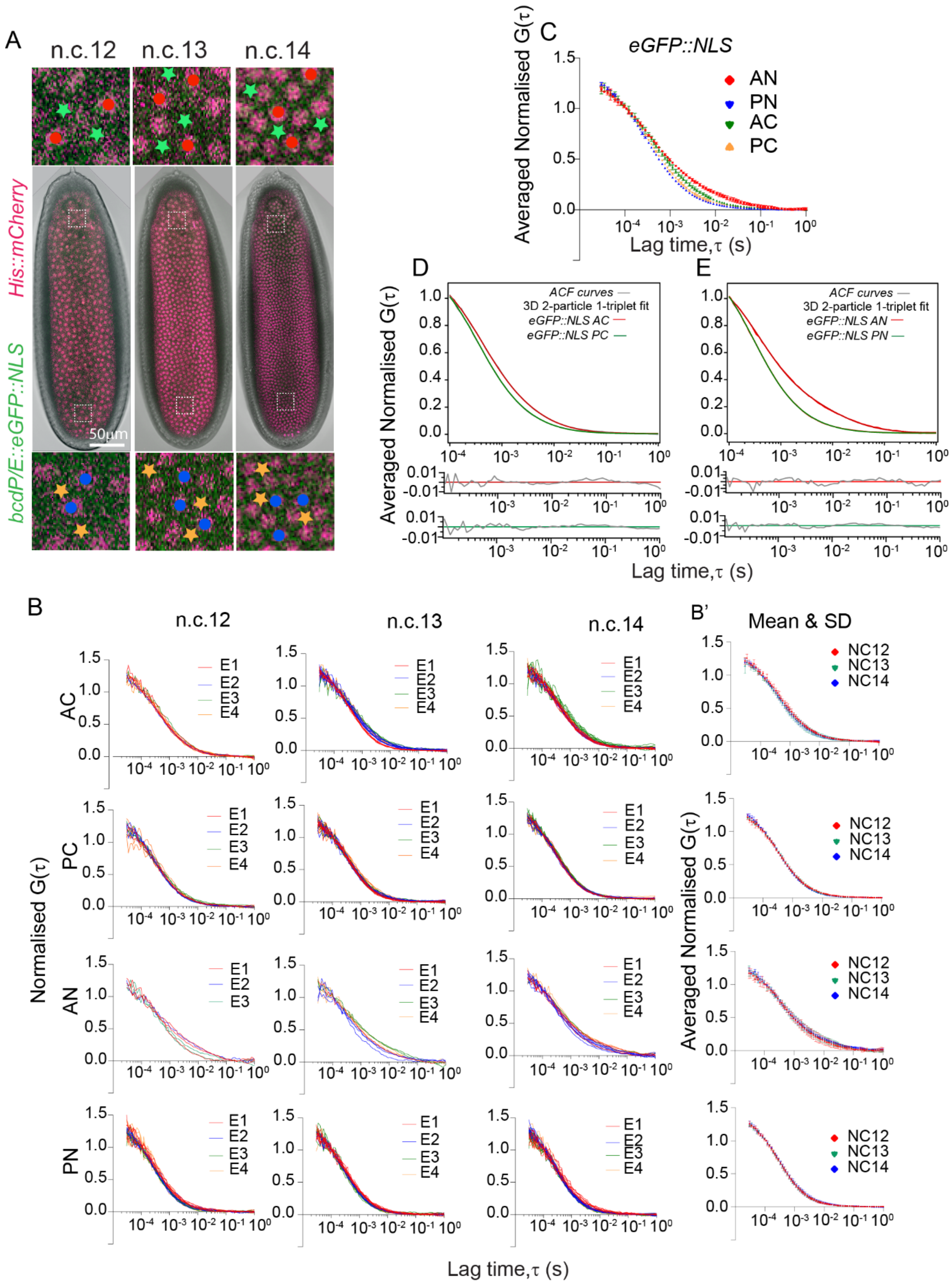


**Figure S2: FCS and fitting of eGFP::NLS (related to Figure 2)**

(A) *Drosophila* blastoderm showing the interphase periods of n.c. 12, 13 and 14. Nuclei (mCherry::His2Av, red) and eGFP::NLS (green). Dots and stars indicate cytoplasmic and nuclear regions, respectively, where FCS measurements are carried out in the anterior (red) and posterior (green). (B-B') Normalised ACF curves with mean and S.D. of eGFP::NLS in cytoplasmic and nuclear compartments of the n.c. 12,13, and 14 interphases. (C) Comparison of normalised, averaged ACF curves with mean and S.D. of eGFP::NLS in cytoplasmic and nuclear compartments in n.c. 12,13, and 14 interphases. (D-E) ACF curves (grey) with residues fitted with 3D 2-particle 1-triplet diffusion model for cytoplasm (D) and nuclei (E).

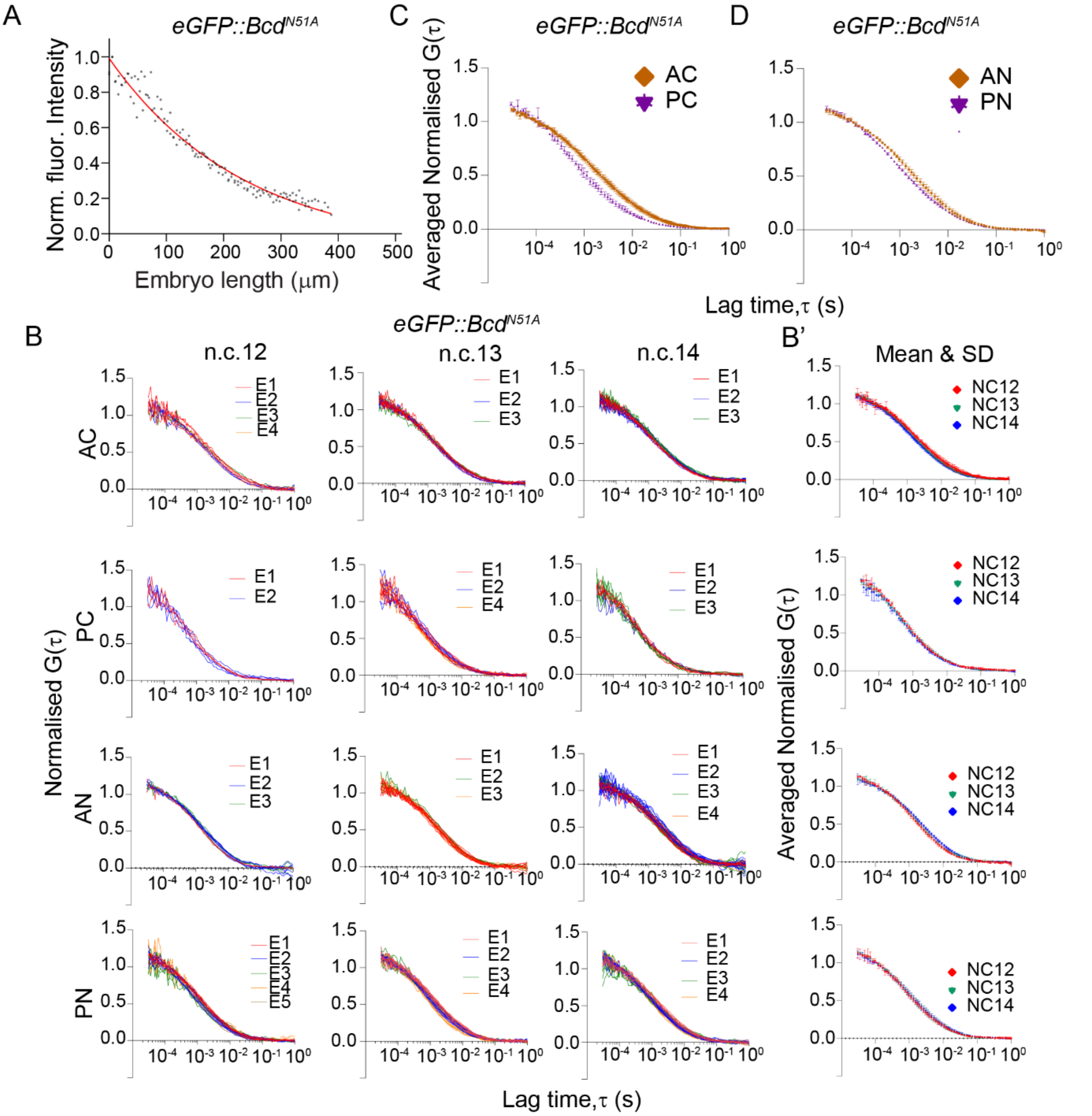


**Figure S3: FCS and fitting of eGFP:Bcd$^{N51A}$ (related to Figure 4)**

(A) Gradient profile of eGFP::Bcd$^{N51A}$ plotted with normalised fluorescence intensities in Y-axis. Red line is fit to exponential profile. (B-B') Normalised ACF curves with mean and S.D. of eGFP::Bcd$^{N51A}$ in the cytoplasmic and nuclear compartments of the n.c. 12,13, and 14 interphases. (C-D) Normalised average ACF curves of eGFP::bcd$^{N51A}$ in the cytoplasmic (C) and nuclear (D) locations of the anterior and posterior domains of the embryo.

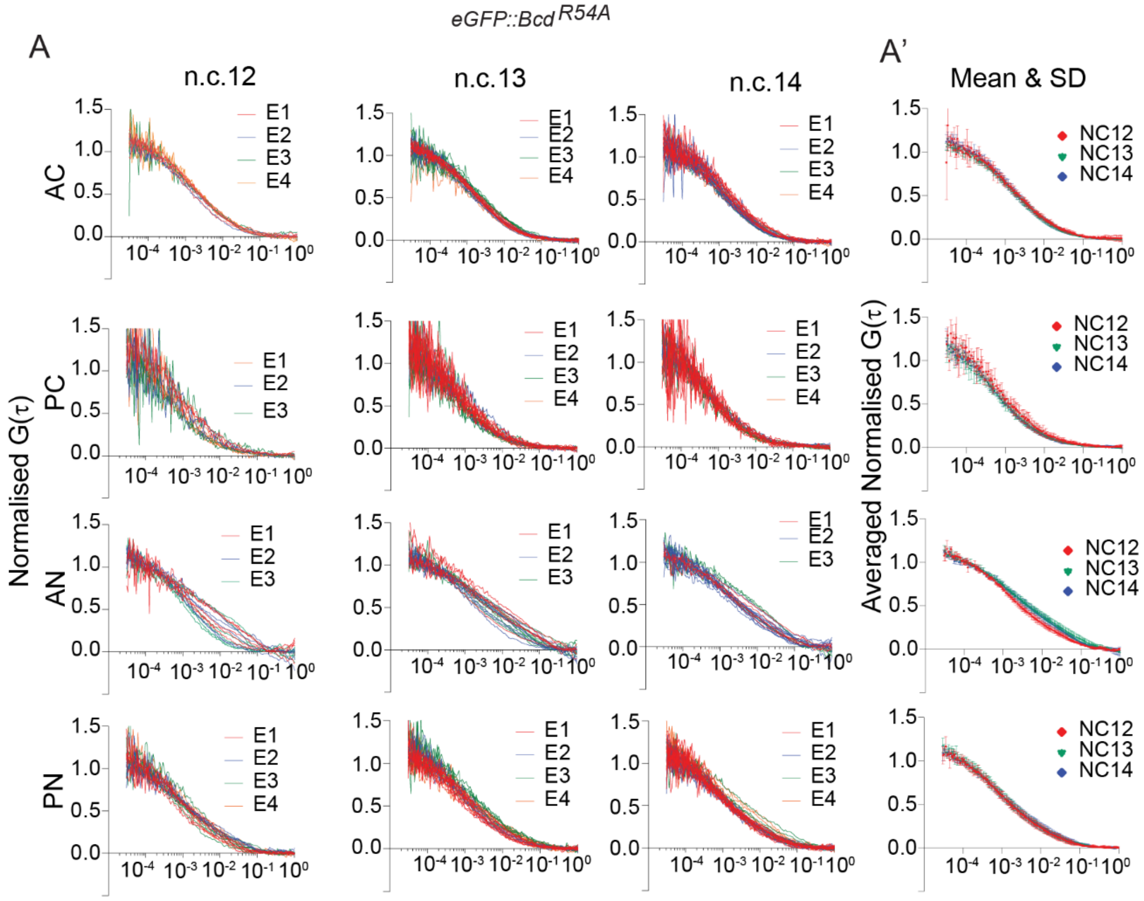


**Figure S4: Dynamics of eGFP::Bcd$^{R54A}$ (related to Figure 4)**

(A) Normalised ACF curves with mean and S.D. of eGFP::Bcd$^{R54A}$ in the cytoplasmic and nuclear compartments of the n.c. 12,13, and 14 interphases. Normalised ACF curves from multiple embryos are shown. Lag times from $10^{-4}$ sec to 1sec are shown for visual clarity. (A') Normalised and averaged autocorrelation ACF curves with mean and S.D. of eGFP::Bcd$^{R54A}$ in the cytoplasmic and nuclear compartments of the n.c. 12,13, and 14 interphases.

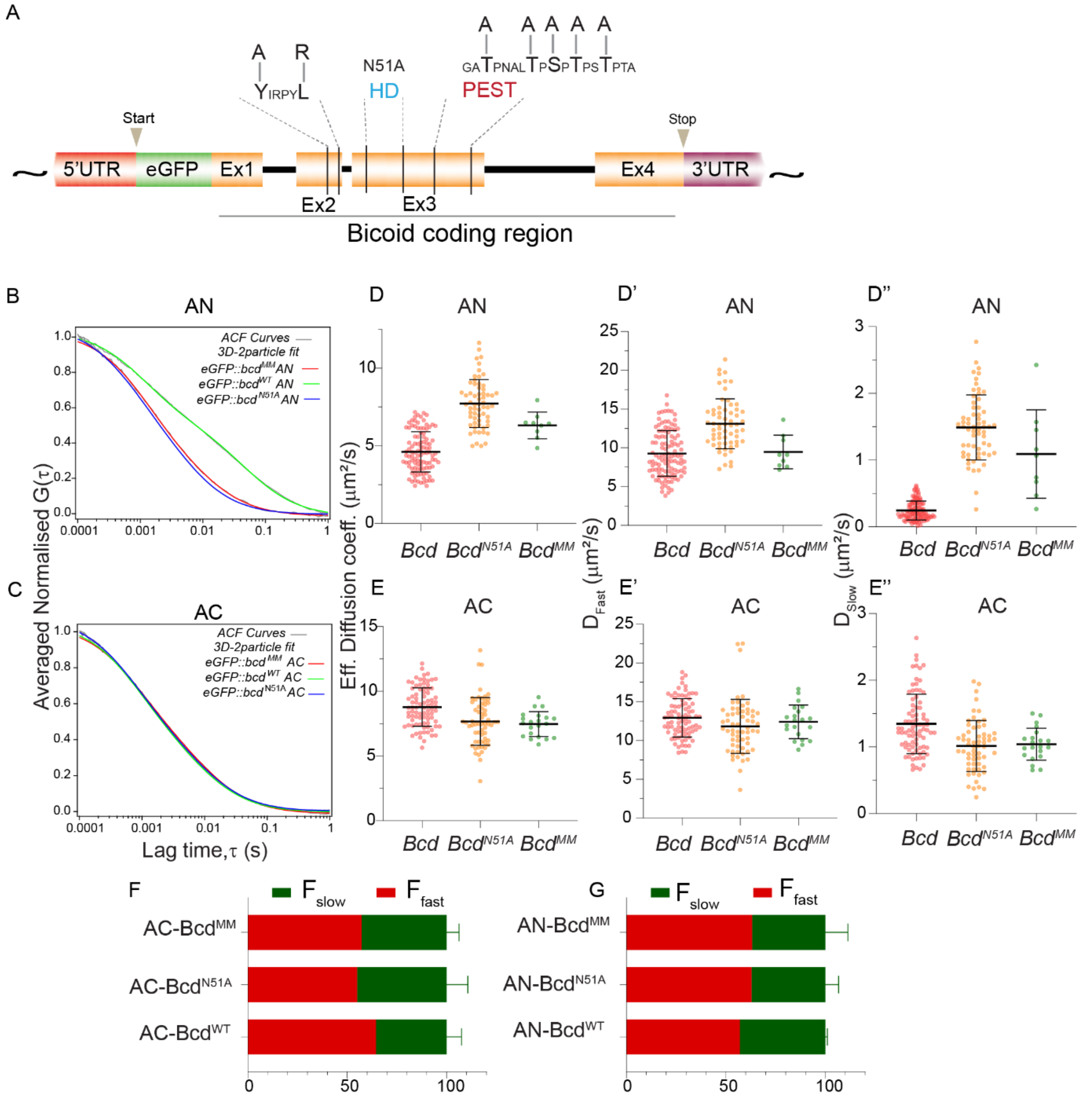


**Figure S5: FCS of eGFP:: Bcd$^{MM}$ (related to Figure 4)**

(A) Schematic of the point mutations introduced in homeodomain, YIRPYL motif and PEST domain that presumably abolishes the function of these domain in eGFP::Bcd$^{MM}$.. (B-C) Normalised ACF curves of eGFP:Bcd, eGFP:bcd$^{N51A}$ and eGFP:Bcd$^{MM}$ from multiple embryos are compared. Lag times from $10^{-4}$ sec to 1sec are shown for visual clarity. (D-E'') Scatter plots of the effective diffusion (D,E), $D_{fast}$ (D',E') and $D_{slow}$ (D''-E'') values compared among eGFP:Bcd, eGFP:Bcd$^{N51A}$ and eGFP:Bcd$^{MM}$ . (F-G) Bar plots comparing the fractions of slow and fast components of the genotypes in D to E''.

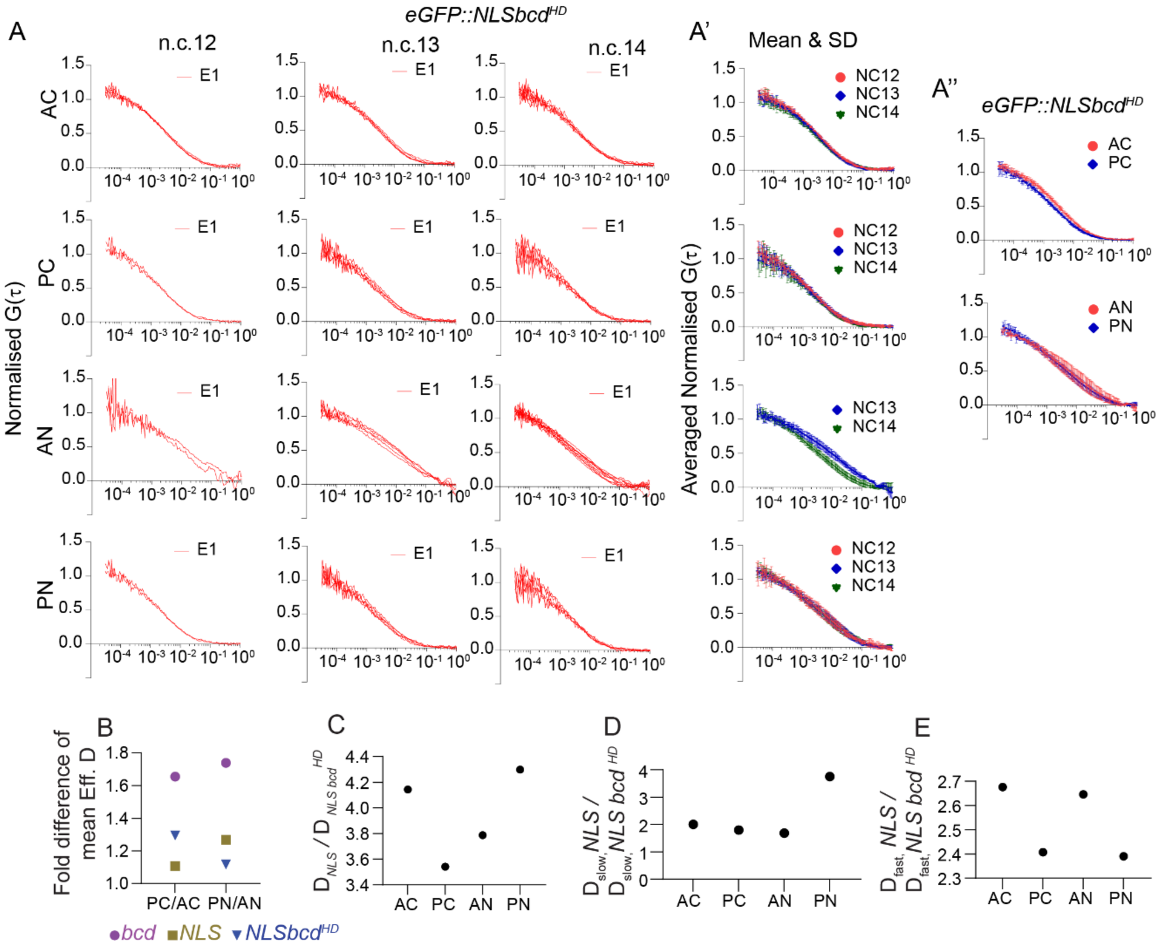


**Figure S6: FCS of eGFP::NLSbcd$^{HD}$ (related to Figure 5)**

(A-A') Normalised ACF curves with mean and S.D. of eGFP::NLSbcd$^{HD}$ in the cytoplasmic and nuclear compartments of the n.c. 12,13, and 14 interphases. Normalised ACF curves from multiple embryos (right) are compared. Lag times from $10^{-4}$ sec to 1sec are shown for visual clarity. (A'') Qualitative comparison of normalised ACF curves with mean and S.D. of eGFP::NLSbcd$^{HD}$ in the cytoplasmic and nuclear compartments of the n.c. 12,13, and 14 interphases. (B-E) Change in relative diffusivity of eGFP::NLSbcd$^{HD}$ compared with eGFP::NLS at different locations within the embryo.

### Supplementary Tables

| N.C. | Embryo No. (no. of ACF curves) | $TauD_1$ (ms) | D1 ($\mu m^2/s$) | Fraction ($F_1\%$) | $TauD_2$ (ms) | $D_2$ ($\mu m^2/s$) | Fraction ($F_2\%$) | Eff.Diff. ($\mu m^2/s$) |
|---|---|---|---|---|---|---|---|---|
| **eGFP::Bcd AC : 3D-Diffusion 2-particle model** | | | | | | | | |
| 12 | 1(4),2(4),3(6),4(4) | 0.8±0.1 | 13.2±1.1 | 66±4 | 8.5±1.7 | 1.3±0.2 | 34±4 | 9.1±0.8 |
| 13 | 1(9),2(8),3(4),4(5) | 0.8±0.1 | 12.8±0.8 | 66±5 | 10.0±1.4 | 1.2±0.2 | 34±3 | 8.7±1.1 |
| 14 | 1(15),2(12),3(12),4(7) | 0.8±0.1 | 12.9±1.1 | 64±5 | 7.9±2.1 | 1.4±0.3 | 36±5 | 8.7±1.1 |
| **eGFP::Bcd PC : 3D-Diffusion 2-particle model** | | | | | | | | |
| 12 | 1(3),2(3),3(3),4(4) | 0.5±0.0 | 19.2±1.6 | 78±3 | 8.5±1.7 | 1.3±0.2 | 22±3 | 15.3±1.6 |
| 13 | 1(4),2(6),3(7),4(3) | 0.6±0.0 | 18.4±0.7 | 77±5 | 7.7±1.3 | 1.3±0.2 | 23±5 | 14.4±0.9 |
| 14 | 1(7),2(7),3(4),4(4) | 0.6±0.0 | 19.0±1.2 | 79±3 | 9.8±1.6 | 1.2±0.1 | 21±3 | 15.2±1.2 |
| **eGFP::NLS AC : 3D-Diffusion 2-particle model** | | | | | | | | |
| 12 | 1(4),2(2),3(3),4(2) | 0.4±0.0 | 25.5±1.7 | 85±2 | 6.8±1.5 | 1.7±0.2 | 15±2 | 21.8±0.9 |
| 13 | 1(4),2(4),3(7),2(4) | 0.4±0.0 | 27.9±3.6 | 84±6 | 6.4±2.6 | 2.0±0.6 | 16±6 | 22.8±1.0 |
| 14 | 1(8),2(7),3(5),4(6) | 0.4±0.0 | 26.4±1.2 | 84±7 | 5.6±2.1 | 2.3±0.5 | 16±7 | 22.5±0.8 |
| **eGFP::NLS PC : 3D-Diffusion 2-particle model** | | | | | | | | |
| 12 | 1(3),2(4),3(3),4(1) | 0.3±0.0 | 28.1±3.2 | 91±0 | 6.4±1.7 | 2.1±1.3 | 9±0 | 26.1±2.6 |
| 13 | 1(3),2(4),3(6),4(5) | 0.4±0.0 | 27.0±1.3 | 92±2 | 5.3±1.8 | 2.5±0.4 | 8±2 | 25.0±1.4 |
| 14 | 1(1),2(4),3(4),4(6) | 0.4±0.0 | 26.4±2.6 | 94±1 | 6.9±2.8 | 2.3±0.6 | 6±1 | 25.1±2.2 |

**Table 1: Comparison of parameter values of cytoplasmic eGFP:Bcd and eGFP:NLS diffusion fit using 3D 2-particle diffusion model.**

| N.C. | Embryo No. (no. of ACF curves) | TauD1 (ms) | D1 ($\mu m^2/s$) | Fraction ($F_1\%$) | TauD2 (ms) | D2 ($\mu m^2/s$) | Fraction ($F_2\%$) | Eff.Diff. ($\mu m^2/s$) |
|---|---|---|---|---|---|---|---|---|
| **eGFP::Bcd AN : 3D-Diffusion 2-particle model** | | | | | | | | |
| 12 | 1(6),2(8),3(5),4(3) | 1.1±0.1 | 9.7±1.3 | 50±4 | 43.0±6.2 | 0.3±0.1 | 50±4 | 4.9±0.5 |
| 13 | 1(9),2(11),3(6),4(8) | 1.2±0.1 | 9.3±0.5 | 49±3 | 66.4±15.6 | 0.3±0.0 | 51±3 | 4.6±0.5 |
| 14 | 1(13),2(10),3(11),4(10) | 1.2±0.1 | 9.0±0.8 | 49±4 | 65.2±7.6 | 0.2±0.0 | 51±4 | 4.5±0.5 |
| **eGFP::Bcd PN : 3D-Diffusion 2-particle model** | | | | | | | | |
| 12 | 1(8), 2(4), 3(6), 4(5) | 0.9±0.1 | 11.9±1.5 | 64±3 | 36.0±9.4 | 0.4±0.1 | 36±3 | 7.7±1.1 |
| 13 | 1(9), 2(9), 3(10), 4(5) | 0.8±0.1 | 13.1±1.3 | 61±1 | 38.3±5 | 0.4±0.1 | 39±1 | 8.1±0.9 |
| 14 | 1(5), 2(8), 3(9), 4(7) | 0.9±0.1 | 11.9±1.3 | 63±2 | 42.2±11 | 0.3±0.1 | 37±2 | 7.5±0.7 |
| **eGFP:NLS AN : 3D-Diffusion 2-particle model** | | | | | | | | |
| 12 | 1(2),2(1),3(2) | 0.4±0.0 | 26.1±0.5 | 84±0 | 6.6±3.3 | 1.7±0.8 | 16±0 | 21.8±0.5 |
| 13 | 1(1),2(4),3(2),4(3) | 0.4±0.0 | 26.1±2.6 | 79±6 | 12.2±2.3 | 0.9±0.3 | 21±6 | 20.4±1.2 |
| 14 | 1(3),2(6),3(1),4(4) | 0.4±0.0 | 27.1±2.0 | 80±5 | 15.7±5.4 | 0.9±0.2 | 20±5 | 21.7±2.6 |
| **eGFP::NLS PN : 3D-Diffusion 2-particle model** | | | | | | | | |
| 12 | 1(5),2(3),3(7),4(2) | 0.3±0.0 | 28.2±2.2 | 95±1 | 9.6±5.0 | 2.1±1.4 | 5±1 | 26.9±2.2 |
| 13 | 1(4),2(5),3(3),4(8) | 0.3±0.0 | 30.3±2.8 | 95±1 | 15.5±9.9 | 1.2±0.3 | 5±1 | 28.9±3.0 |
| 14 | 1(4),2(8),3(7),4(6) | 0.3±0.0 | 28.3±2.8 | 95±3 | 14.8±8.3 | 1.3±0.5 | 5±3 | 26.9±2.9 |

**Table 2: Comparison of parameter values of nuclear eGFP:Bcd and eGFP:NLS diffusion fit using 3D 2-particle diffusion model.**

| N.C. | Embryo No. (no. of ACF curves) | $TauD_1$ (ms) | D1 ($\mu m^2/s$) | Fraction ($F_1$%) | $TauD_2$ (ms) | $D_2$ ($\mu m^2/s$) | Fraction ($F_2$%) | Eff.Diff. ($\mu m^2/s$) |
|---|---|---|---|---|---|---|---|---|
| **eGFP::Bcd$^{N51A}$ AN: 3D-Diffusion 2-particle model** | | | | | | | | |
| 12 | 1(3),2(6),3(4) | 0.7±0.1 | 16.1±3.8 | 53±3 | 5.3±0.9 | 2.0±0.4 | 47±3 | 9.3±1.5 |
| 13 | 1(6),2(6),4(3) | 0.8±0.1 | 13.5±2.4 | 56±1 | 6.6±0.7 | 1.6±0.2 | 44±1 | 8.1±1.7 |
| 14 | 1(11),2(6),3(8),4(10) | 0.8±0.0 | 13.0±1.0 | 52±1 | 8.0±1.2 | 1.4±0.1 | 48±1 | 7.4±0.7 |
| **eGFP::Bcd$^{N51A}$ PN: 3D-Diffusion 2-particle model** | | | | | | | | |
| 12 | 1(6),2(7),3(3),4(3),5(4) | 0.7±0.0 | 15.8±1.5 | 69±5 | 7.0±1.2 | 1.7±0.4 | 31±5 | 11.0±1.3 |
| 13 | 1(11),2(10),3(5), 4(5),5(6) | 0.7±0.1 | 16.0±2.2 | 67±5 | 7.0±1.8 | 1.5±0.2 | 33±5 | 11.0±2.0 |
| 14 | 1(7),2(12),3(8),4(6),5(9) | 0.6±0.1 | 17.0±3.2 | 64±6 | 7.0±1.2 | 1.6±0.2 | 36±6 | 11.0±2.0 |
| **eGFP::Bcd$^{N51A}$ AC: 3D-Diffusion 2-particle model** | | | | | | | | |
| 12 | 1(4),2(2),3(1),4(2) | 1.0±0.2 | 10.7±2.7 | 64±6 | 14.1±4.3 | 0.8±0.3 | 36±6 | 7.1±1.3 |
| 13 | 1(6),2(6),3(5),4(6) | 1.0±0.3 | 10.7±2.4 | 64±3 | 13.4±4.3 | 0.9±0.3 | 36±3 | 7.0±1.4 |
| 14 | 1(6),2(7),3(8),4(8) | 0.9±0.3 | 12.3±2.7 | 64±2 | 11.8±5.4 | 1.0±0.3 | 36±2 | 8.0±1.7 |
| **eGFP::Bcd$^{N51A}$ PC: 3D-Diffusion 2-particle model** | | | | | | | | |
| 12 | 1(4),2(2),3(4) | 0.6±0.1 | 16.9±0.6 | 79±7 | 9.5±1.7 | 1.2±0.3 | 20±7 | 13.5±1.6 |
| 13 | 1(6),2(6),3(4),4(8),5(7) | 0.6±0.0 | 18.5±2.8 | 79±3 | 9.7±2.5 | 1.3±0.5 | 21±3 | 14.7±2.2 |
| 14 | 1(6),2(4),3(6),4(6) | 0.6±0.0 | 18.3±1.1 | 83±6 | 11.3±1.7 | 1.1±0.3 | 17±6 | 15.2±1.2 |
| **eGFP::Bcd$^{R54A}$ AN: 3D-Diffusion 2-particle model** | | | | | | | | |
| 12 | 1(5),2(2),3(5) | 1.2±0.2 | 9.3±0.4 | 59±3 | 37.0±8.0 | 0.4±0.1 | 41±3 | 5.5±0.0 |
| 13 | 1(5),2(4),3(9) | 1.4±0.2 | 8.1±1.3 | 55±1 | 57.8±15.0 | 0.3±0.1 | 45±1 | 4.5±0.7 |
| 14 | 1(9),2(12),3(7) | 1.3±0.2 | 8.5±1.6 | 55±2 | 40.1±10.0 | 0.4±0.1 | 45±2 | 4.8±1.1 |
| **eGFP::Bcd$^{R54A}$ PN: 3D-Diffusion 2-particle model** | | | | | | | | |
| 12 | 1(5),2(3),3(7) | 0.7±0.1 | 14.4±3.0 | 70±5 | 19.0±5.8 | 0.7±0.2 | 30±5 | 10.2±1.9 |
| 13 | 1(9),2(5),3(7) | 0.7±0.0 | 15.2±1.0 | 70±2 | 16.0±1.0 | 0.7±0.1 | 30±2 | 10.7±0.7 |
| 14 | 1(13),2(8),3(9) | 0.7±0.0 | 16.0±1.0 | 69±1 | 18.0±4.0 | 0.7±0.1 | 31±1 | 11.1±0.6 |
| **eGFP::Bcd$^{R54A}$ AC: 3D-Diffusion 2-particle model** | | | | | | | | |
| 12 | 1(5),2(3),3(2),4(2) | 0.9±0.1 | 11.4±1.7 | 66±4 | 10.5±1.6 | 1.0±0.1 | 34±4 | 7.8±1.2 |
| 13 | 1(10),2(7),3(8),4(5) | 1.0±0.2 | 11.3±1.8 | 66±3 | 10.0±3.0 | 1.1±0.3 | 34±3 | 7.7±1.3 |
| 14 | 1(10),2(13),3(3),4(11) | 0.9±0.1 | 11.8±0.7 | 67±7 | 9.7±2.0 | 1.2±0.2 | 33±7 | 8.2±0.2 |
| **eGFP::Bcd$^{R54A}$ PC: 3D-Diffusion 2-particle model** | | | | | | | | |
| 12 | 1(4),2(4),3(3),4(4) | 0.8±0.2 | 13.8±2.4 | 84±6 | 21.3±5.0 | 0.6±0.1 | 16±6 | 12.5±1.5 |
| 13 | 1(10),2(3),3(8),4(7) | 0.8±0.1 | 13.8±2.6 | 86±2 | 22.0±5.9 | 0.7±0.2 | 14±2 | 11.8±2.3 |
| 14 | 1(7),2(3),3(7),4(4) | 0.6±0.1 | 17.6±2.9 | 83±5 | 15.0±10.7 | 1.0±0.5 | 17±5 | 14.2±1.9 |

**Table 3: Comparison of parameter values of eGFP::Bcd$^{N51A}$ and eGFP::Bcd$^{R54A}$ diffusion fit using 3D 2-particle diffusion model.**

| N.C. | Embryo No. (no. of ACF curves) | TauD$_1$ (ms) | D1 ($\mu m^2/s$) | Fraction ($F_1$%) | TauD$_2$ (ms) | D$_2$ ($\mu m^2/s$) | Fraction ($F_2$%) | Eff.Diff. ($\mu m^2/s$) |
|---|---|---|---|---|---|---|---|---|
| **eGFP::NLSbcd$^{HD}$ AN : 3D-Diffusion 2-particle model** | | | | | | | | |
| 12 | 1(2) | 2.3±0.1 | 4.3±0.2 | 49±1 | 55.6±33.3 | 0.2±0.1 | 51±1 | 2.2±0.2 |
| 13 | 1(10) | 1.2±0.3 | 9.0±2.2 | 52±13 | 64.3±53.8 | 0.3±0.3 | 48±13 | 4.7±1.5 |
| 14 | 1(12) | 0.8±0.3 | 13.3±4.3 | 53±11 | 19.0±8.8 | 0.6±0.3 | 47±11 | 7.2±2.2 |
| **eGFP::NLSbcd$^{HD}$ PN : 3D-Diffusion 2-particle model** | | | | | | | | |
| 12 | 1(5) | 0.7±0.1 | 14.8±2.7 | 44±13 | 19.3±5.9 | 0.6±0.2 | 56±13 | 6.7±1.2 |
| 13 | 1(10) | 0.8±0.3 | 14.0±4.6 | 51±16 | 18.2±6.1 | 0.6±0.2 | 49±16 | 7.3±2.6 |
| 14 | 1(7) | 1.3±0.3 | 8.5±2.4 | 54±7 | 25.2±9.8 | 0.4±0.0 | 46±7 | 5.1±0.2 |
| **eGFP::NLSbcd$^{HD}$ AC : 3D-Diffusion 2-particle model** | | | | | | | | |
| 12 | 1(4) | 0.4±0.0 | 26.1±0.5 | 46±13 | 8.5±2.4 | 1.3±0.4 | 54±12 | 5.3±0.3 |
| 13 | 1(7) | 0.4±0.0 | 26.1±2.6 | 39±8 | 7.3±2.3 | 1.4±0.3 | 61±8 | 6.2±1.6 |
| 14 | 1(7) | 0.3±0.0 | 28.1±0.1 | 50±5 | 8.2±0.9 | 1.2±0.1 | 50±4 | 6.0±1.0 |
| **eGFP::NLSbcd$^{HD}$ PC : 3D-Diffusion 2-particle model** | | | | | | | | |
| 12 | 1(3) | 0.3±0.0 | 28.1±3.2 | 57±0 | 7.8±1.4 | 1.3±0.2 | 43±0 | 6.7±0.4 |
| 13 | 1(6) | 0.3±0.0 | 27.0±1.3 | 59±7 | 12.6±9.1 | 1.0±0.5 | 41±7 | 6.4±1.0 |
| 14 | 1(9) | 0.3±0.0 | 26.4±2.6 | 59±10 | 9.2±3.7 | 1.2±0.4 | 41±10 | 7.2±2.2 |

**Table 4: Comparison of parameter values of eGFP::NLSbcd$^{HD}$ diffusion fit using 3D -2-particle diffusion model.**
